# BDNF controls bidirectional endocannabinoid-plasticity at corticostriatal synapses

**DOI:** 10.1101/550947

**Authors:** Giuseppe Gangarossa, Sylvie Perez, Yulia Dembitskaya, Ilya Prokin, Hugues Berry, Laurent Venance

## Abstract

The dorsal striatum exhibits bidirectional corticostriatal synaptic plasticity, NMDAR- and endocannabinoids-(eCB)-mediated, necessary for the encoding of procedural learning. Therefore, characterizing factors controlling corticostriatal plasticity is of crucial importance. Brain-derived neurotrophic factor (BDNF) and its receptor, the tropomyosine receptor kinase-B (TrkB), shape striatal functions and their dysfunction deeply affects basal ganglia. BDNF/TrkB signaling controls NMDAR-plasticity in various brain structures including the striatum. However, despite cross-talk between BDNF and eCBs, the role of BDNF in eCB-plasticity remains unknown. Here, we show that BDNF/TrkB signaling promotes eCB-plasticity (LTD and LTP) induced by rate-based (low-frequency stimulation) or spike-timing-based (spike-timing-dependent plasticity, STDP) paradigm in striatum. We show that TrkB activation is required for the expression and the scaling of both eCB-LTD and eCB-LTP. Using two-photon imaging of dendritic spines combined with patch-clamp recordings, we show that TrkB activation prolongs intracellular calcium transients, thus increasing eCB synthesis and release. We provide a mathematical model for the dynamics of the signaling pathways involved in corticostriatal plasticity. Finally, we show that TrkB activation enlarges the domain of expression of eCB-STDP. Our results reveal a novel role for BDNF/TrkB signaling in governing eCB-plasticity expression in striatum, and thus the engram of procedural learning.

## INTRODUCTION

The striatum, the main input structure of the basal ganglia system, plays a pivotal role in action selection and procedural learning (Graybiel and Grafton, 2015). Long-term synaptic plasticity at corticostriatal synapses has been associated with the acquisition of a complex and dynamic behavioral repertoire (Yin et al., 2009; Koralek et al., 2012; Shan et al., 2014; Rothwell et al., 2015; Hawes et al., 2015; Xiong et al., 2015; Ma et al., 2018; reviewed in Perrin and Venance, 2018). Upon various activity-dependent patterns (rate-or spike-timing-coded), corticostriatal synapses exhibit bidirectional plasticity, LTP and LTD, mediated by NMDA and type-1 cannabinoid receptors (CB1Rs), respectively (Di Filippo et al., 2009; Lovinger, 2010). Thus, identifying factors modulating the expression of activity-dependent plasticity (Foncelle et al., 2018) in the striatum is of crucial importance.

Emerging evidences suggest that the brain-derived neurotrophic factor (BDNF) and its receptor, the tropomyosine receptor kinase B (TrkB) (Deinhardt and Chao, 2014), are strongly involved in shaping striatal functions (Baydyuk et al., 2011; Besusso et al., 2013; Unterwald et al., 2013; Jing et al., 2017). In addition, alterations of BDNF/TrkB signaling have been observed in several psychiatric and neurodegenerative disorders (Zuccato and Cattaneo, 2009; Autry and Monteggia, 2012). The involvement of BDNF in the expression of (striatal) NMDAR-mediated LTP is now well elucidated (Jia et al., 2010; Park and Poo, 2013; Park et al., 2014). Despite multiple reports revealing functional cross-talk between BDNF and endocannabinoids (eCBs) signaling in the cortex (Huang et al., 2008; Lemtiri-Chlieh and Levine, 2010; Yeh et al., 2017; Maglio et al., 2018), hippocampus (Khaspekov et al., 2004; Roloff et al., 2010), ventral tegmental area (Zhong et al., 2015) and cerebellum (Maison et al., 2009), the role of BDNF in eCB-mediated long-term synaptic plasticity remains largely unknown. Here, we asked whether BDNF/TrkB signaling exerts a control over eCB-plasticity at corticostriatal synapses in the dorsal striatum.

eCBs have emerged as a major signaling system in learning and memory because of their involvement in synaptic plasticity (Castillo et al., 2012; Araque et al., 2017). The eCB system comprises active lipids (mainly 2-arachidonylglycerol, 2-AG, and anandamide) synthesized and released on-demand, which act as retrograde neurotransmitters on presynaptic type-1 cannabinoid receptors (CB1Rs) and postsynaptic transient receptor potential vanilloid-type-1 (TRPV1) (Piomelli et al., 2007; Alger and Kim, 2011). In the dorsal striatum very low levels of BDNF have been reported and the striatal output neurons, the medium-sized spiny neurons (MSNs), do not show detectable level of BDNF mRNA (Altar et al., 1997; Conner et al., 1997). However, MSNs display high levels of TrkB (Baydyuk et al., 2011; Besusso et al., 2013; Unterwald et al., 2013). Striatal BDNF is mainly released from glutamatergic cortical afferents (Altar et al., 1997; Jia et al., 2010). Here, we took advantage of various forms of activity-dependent eCB-plasticity at corticostriatal synapses to investigate the role of BDNF/TrkB signaling in these plasticities. Indeed, in the dorsal striatum, at least three forms of eCB-plasticity have been observed: (*i*) an eCB-LTD induced by a low frequency stimulation (LFS) (Fino et al., 2005; Puente et al., 2011), (*ii*) an eCB-LTD induced by 100-150 spike-timing-dependent plasticity (STDP) (eCB-tLTD) (Pawlak and Kerr, 2008; Shen et al., 2008; Fino et al., 2010; Paillé et al., 2013), and (*iii*) a eCB-tLTP induced by low numbers (~10-15) of STDP pairings (Cui et al., 2015; Cui et al., 2016; Cui et al., 2018; Xu et al., 2018).

Considering the cross-talk between BDNF/TrkB and eCB signaling in promoting eCB synthesis (Zhao et al., 2015; Bennett et al., 2017), we questioned the role of TrkB in the expression of eCB-plasticity at corticostriatal synapses. Here, we provide evidence for a novel role of BDNF/TrkB in promoting eCB-mediated synaptic plasticity induced by rate-based or spike-timing-based paradigm. We show that TrkB activation is required for the expression and the scaling of both eCB-LTD and eCB-LTP at the glutamatergic corticostriatal synapse. Using two-photon imaging at the level of the dendritic elements (spines and shafts) of MSNs combined with patch-clamp recording, our results show that TrkB activation acts as a molecular trigger for boosting intracellular Ca^2+^ transients, thus increasing eCB synthesis and release. We also provide a mathematical model for the dynamics of the signaling pathways involved in striatal plasticity. Combining our experimental results and mathematical modeling, we show that TrkB activation is not only required for eCB-plasticity expression but also for the scaling of the domain of eCB-plasticity expression, in particular in STDP paradigms. Our results reveal a novel role for BDNF/TrkB as a key factor for the control of the Hebbian eCB-plasticity map in the dorsal striatum.

## MATERIALS AND METHODS

### Animals

All experiments were performed in accordance with the guidelines of the local animal welfare committee (Center for Interdisciplinary Research in Biology Ethics Committee) and the EU (directive 2010/63/EU). Every precaution was taken to minimize stress and the number of animals used in each series of experiments. Pre-adult OFA rats P_25-35_ (Charles River, L’Arbresle, France) were used for brain slice electrophysiology. Animals were housed in standard 12-hour light/dark cycles and food and water were available *ad libitum*.

### Brain slice preparation and patch-clamp recordings

Horizontal brain slices (300-330 µm-thick) containing the somatosensory cortical area and the corresponding corticostriatal projection field (Fino et al., 2005; Cui et al., 2015) were prepared with a vibrating blade microtome (VT1200S, Leica Microsystems, Nussloch, Germany). Corticostriatal connections (between somatosensory cortex layer 5 and the dorso-lateral striatum) are preserved in the horizontal plane. Brains were sliced in a 95% CO2/5% O2-bubbled, ice-cold cutting solution containing (in mM) 125 NaCl, 2.5 KCl, 25 glucose, 25 NaHCO3, 1.25 NaH2PO4, 2 CaCl2, 1 MgCl2, 1 pyruvic acid, and transferred into the same solution at 34°C for one hour and next moved to room temperature. Whole-cell recordings were performed as previously described (Fino et al., 2010; Paillé et al., 2013). For whole-cell recordings, borosilicate glass pipettes of 6-8MΩ resistance were filled with (in mM): 105 K-gluconate, 30 KCl, 10 HEPES, 10 phosphocreatine, 4 Mg-ATP, 0.3 Na-GTP, 0.3 EGTA (adjusted to pH 7.35 with KOH). The composition of the extracellular solution was (mM): 125 NaCl, 2.5 KCl, 25 glucose, 25 NaHCO_3_, 1.25 NaH_2_PO_4_, 2 CaCl_2_, 1 MgCl_2_, 10 µM pyruvic acid bubbled with 95% O_2_ and 5% CO_2_. Signals were amplified using EPC10-2 amplifiers (HEKA Elektronik, Lambrecht, Germany). All recordings were performed at 34°C, using a temperature control system (Bath-controller V, Luigs & Neumann, Ratingen, Germany) and slices were continuously superfused with extracellular solution, at a rate of 2 ml/min. Slices were visualized under an Olympus BX51WI microscope (Olympus, Rungis, France), with a 4x/0.13 objective for the placement of the stimulating electrode and a 40x/0.80 water-immersion objective for the localization of cells for whole-cell recordings. Current- and voltage-clamp recordings were sampled at 10 kHz, with the Patchmaster v2×32 program (HEKA Elektronik).

### Synaptic plasticity induction protocols

Electrical stimulations were performed with a concentric bipolar electrode (Phymep, Paris, France) placed in layer 5 of the somatosensory cortex. Electrical stimulations were monophasic, at constant current (ISO-Flex stimulator, AMPI, Jerusalem, Israel). Currents were adjusted to evoke 100-300 pA EPSCs. Repetitive control stimuli were applied at 0.1 Hz.

#### Low frequency stimulation protocols

Low-frequency stimulation (LFS) consisted in 600 cortical stimuli at 1 Hz and was performed in a Hebbian mode; Indeed, the depolarization of the postsynaptic element from its resting membrane potential (RMP) to 0 mV was coincident with the presynaptic stimulation.

#### Spike-timing-dependent plasticity protocols

STDP protocols consisted of pairings of pre- and postsynaptic stimulations (at 1 Hz) separated by a specific time interval (Δt_STDP_). Presynaptic stimulations corresponded to cortical stimulations and the postsynaptic stimulation of an action potential evoked by a depolarizing current step (30 ms duration) in MSNs. The STDP protocol involved pairing pre- and postsynaptic stimulation with a certain fixed timing interval, Δt_STDP_ (Δt_STDP_<0 indicating that postsynaptic stimulation preceded presynaptic stimulation, i.e. post-pre pairings, and Δt_STDP_>0 indicating that presynaptic stimulation preceded postsynaptic stimulation, i.e. pre-post pairings), repeated n times at 1 Hz. eCB-tLTD was induced with 100 pre-post pairings with 10<Δt_STDP_<20 ms and eCB-tLTP was induced with 10 post-pre pairings with −20<Δt_STDP_<-10 ms as previously described (Cui et al., 2015; Cui et al., 2016; Xu et al., 2018). Recordings were made over a period of 10 minutes at baseline, and for at least 40 minutes after the STDP induction protocols; long-term changes in synaptic efficacy were measured for the last 10 minutes. We individually measured and averaged 60 successive EPSCs, comparing the last 10 minutes of the recording with the 10-minute baseline recording. Neuron recordings were made in voltage-clamp mode during baseline and for the 50-60 minutes of recording after the STDP protocol. Variation of input and access resistances, measured every 10 sec all along the experiment, beyond 20% led to the rejection of the experiment.

### Electrophysiological data analysis

Off-line analysis was performed with Fitmaster (Heka Elektronik), Igor-Pro 6.0.3 (Wavemetrics, Lake Oswego, OR, USA) and custom-made software in Python 3.0. Statistical analysis was performed with Prism 5.02 software (San Diego, CA, USA). In all cases “n” refers to an experiment on a single cell from a single slice. All results are expressed as mean ± SEM. Statistical significance was assessed by two-tailed student *t*-tests (unpaired or paired *t*-tests) or one-way ANOVA (with Newman-Keuls post hoc test) when appropriate, using the indicated significance threshold (*p*).

### Two-photon imaging combined with whole-cell recordings

Whole-cell patch-clamp pipettes (6-8 MΩ) were filled with the solution (in mM): 122 K-gluconate, 13 KCl, 10 HEPES, 10 phosphocreatine, 4 Mg-ATP, 0.3 Na-GTP, 0.3 EGTA (adjusted to pH 7.35 with KOH). Morphological tracer Alexa Fluor 594 (50 µM) and Ca^2+^-sensitive dye Fluo-4F (250 µM) were added to the intracellular solution to monitor Ca^2+^ transients. Cells were filled with the dyes for at least 15 min to ensure dye equally distributed and visually identified under Scientifica TriM Scope II system (LaVision, Germany), with a 60x/1.00 water-immersion objective. Alexa Fluor 594 and Fluo-4F were excited at 830 nm wavelength (femtoseconds IR laser Chameleon MRU-X1, Coherent, UK), and their fluorescence were detected with photomultipliers within 525/50 and 620/60 nm ranges, respectively. Line-scan imaging at 200 Hz was performed to obtain Ca^2+^ signals in the dendritic shaft and spines and was synchronized with patch-clamp recordings. In each recording we injected a prolonged somatic depolarization and monitored maximal Ca^2+^ elevations to verify linear dependence of Fluo-4F Ca^2+^ signals (nonlinearity was below 20%) and that Ca^2+^ transients were below saturation level. The changes in baseline Ca^2+^ level were monitored as the ratio between the baseline Fluo-4F and Alexa Fluor 594 fluorescence. If this ratio increased during the experiment for more than 20%, the cell was discarded. The dark noise of the photomultipliers was collected when the laser shutter was closed in every recording. We chose to examine the τ decay constant of Ca^2+^ response (τCa^2+^) and not the calcium amplitude because of run-down phenomenon.

Two back-propagating action potentials (bAPs) evoked by a depolarizing current step (30 ms duration) have been used to monitor Ca^2+^ transients either in an unpaired manner (i.e. bAPs only) or paired with cortical stimulation (post-pre or pre-post paired stimulations). The excitability was measured in current-clamp mode by 500 ms steps of current injections from - 300 to +500 pA with step of 20 pA to verify the identity of the cell and measure input resistance. Input and series resistances were routinely measured in voltage-clamp mode and data were discarded if resistances changed by more than 20% during the recording. Signals were amplified using with EPC10-2 amplifiers (HEKA Elektronik, Lambrecht, Germany). All recordings were performed at 34°C (Bath-controller-V, Luigs & Neumann, Ratingen, Germany) and slices were continuously superfused with extracellular solution, at a rate of 2 ml/min. Recordings were sampled at 10 kHz with the Patchmaster v2×32 program (HEKA Elektronik).

Electrophysiological data were analyzed with Fitmaster (Heka Elektronik) and custom-made software in Python 3.0. Ca^2+^ transients were analyzed with custom made software in Python 3.0 and averaged in MS Excel (Microsoft, USA). The measurements of Ca^2+^ transient were represented as ΔG/R: (G_peak_-G_baseline_)/(R_baseline_-R_dark noise_). Baseline Ca^2+^ signals were represented by baseline G/R, (G_baseline_-G_dark noise_)/(R_baselin-Rdark noise_), where G is the Fluo-4F fluorescence, and R is Alexa Fluor 594 fluorescence. G_baseline_, R_baseline_ and G_peak_ were obtained from the parameters of the bi-exponential fitting model in each trial and then averaged between 5-6 repetitions for each condition; the bi-exponential fitting was chosen to maximize the fit of the Ca2+-evoked events and gave better fit quality than single- and triple-exponential fitting (https://github.com/calciumfilow/calcium-trace-extractor). G_dark noise_ and R_dark noise_ are the dark currents of the corresponding photomultipliers.

We used ratiometric dyes to monitor Ca^2+^ variations (Yasuda et al., 2004), to improve signal-to-noise ratio and to lower the dependence of changes in baseline fluorescence. We monitored G_baseline_, R_baseline_ and G_baseline_/R_baseline_ during the whole recording and ensured their stability between control recordings and subsequent drug applications. The stability of Alexa Fluor 594 fluorescence was assessed as R_baseline_ after the first (~+35-40 min) and the second (~+50-55 min) drug applications compared to control (~+15-25 min) measurement in dendritic spines and shafts; R_baseline_ did not exceed 20% over recording. The ratio of baseline green fluorescence over red fluorescence (G_baseline_/R_baseline_) is a gauge of cell health (Yasuda et.al., 2004) and was also monitored after the first and the second drug applications. G_baseline_/R_baseline_ was compared to control measurement in dendritic spines and shafts and did not exceed 20% over recording. For illustration purposes, traces result from the average of five sequential traces. The statistical significance was tested using a paired or unpaired Student’s t-test in Prism 5.02 software (San Diego, CA, USA). The data are given in mean ± SEM where “n” designates the number of recordings.

### Chemicals

N-(piperidin-1-yl)-5-(4-iodophenyl)-1-(2,4-dichlorophenyl)-4-methyl-1H-pyrazole-3-carboxamide (AM251, 3µM) (Tocris) was dissolved in ethanol and then added in the external solution at a final concentration of ethanol of 0.015%. N-[2-[[(Hexahydro-2-oxo-1H-azepin-3-yl)amino]carbonyl]phenyl]benzo[b]thiophene-2-carboxamide (ANA12, 10 µM) (Tocris), (9S,10R,12R)-2,3,9,10,11,12-Hexahydro-10-hydroxy-9-methyl-1-oxo-9,12-epoxy-1H-diindolo[1,2,3-fg:3’,2’,1’-kl]pyrrolo[3,4-i][1,6]benzodiazocine-10-carboxylic acid methyl ester (K252a, 200 nM) (Tocris), 7,8-Dihydroxy-2-phenyl-4H-1-benzopyran-4-one (DHF, 10 µM) (Tocris), 1,4-Diamino-2,3-dicyano-1,4-bis(2-aminophenylthio)-butadiene (U0126, 10µM) (Tocris) and 2-(4-morpholino)-8-phenyl-4H-1-benzopyran-4-one (LY294002, 10µM) (Tocris) were dissolved in DMSO and then added in the external solution at a final concentration of DMSO of 0.001-0.04%. DMSO (0.001-0.04% final concentration) was used as control for all the experiments.

None of the bath-applied drugs had a significant effect on basal synaptic transmission. The normalized EPSC_(baseline-drugs/baseline-control)_ amplitude were (in %): 97±4 for K252a (200 nM) (n=14, *p*=0.6790), 101±5 for ANA12 (10 µM) (n=12, *p*=0.8001), 97±2 for DHF (10 µM) (n=11, *p*=0.4937) and 100±1 for U0126 (10 µM) co-applied with LY294002 (10 µM) (n=5, *p*=0.2588).

### Mathematical modeling

To account for TrkB signaling in the postsynaptic neuron, the mathematical model of STDP at corticostriatal synapses previously described (Cui et al, 2016) was extended. We refer to Cui et al. (2016) for a complete description of the model and its parameters, and here we give a broad outline of the model, together with a detailed description of the modifications we implemented in the present study. Cui et al. (2016) considers an isopotential electrical model of the postsynaptic membrane potential coupled to a detailed description of postsynaptic signaling pathways, including calcium currents, the activation of CaMKII α and the production of endocannabinoids (Fig. 5A). Each presynaptic stimulus *i* of the STDP protocol triggers a surge of glutamate concentration in the synaptic cleft, *G*(*t*), modeled as an immediate increase followed by exponential decay:

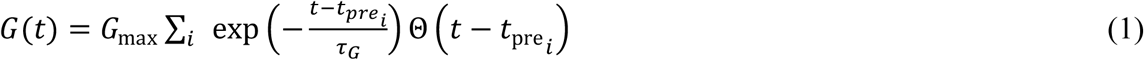

where 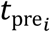 is the time of the presynaptic stimulus and the Heaviside function Θ(*x*) = 1 if *x* ≥ 0, 0 otherwise. Postsynaptic stimulations of the STDP induction protocol give rise to action currents that combine the step-current injected in the postsynaptic soma at time 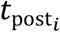 and the resulting action potential:

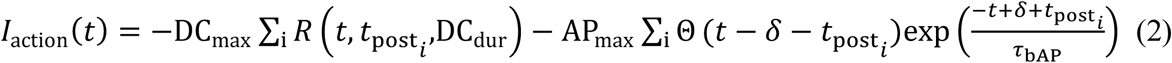

where *R*(*t, a, l*) = Θ(*t* - *a*) - Θ(*t* - *a* - *l*) and *δ* accounts for the time elapsed between the onset of the postsynaptic step current and the action potential (~3ms in MSNs). The model describes the resulting electrical response of an isopotential postsynaptic element endowed with AMPAR, NMDAR, VSCC and TRPV1 conductances:

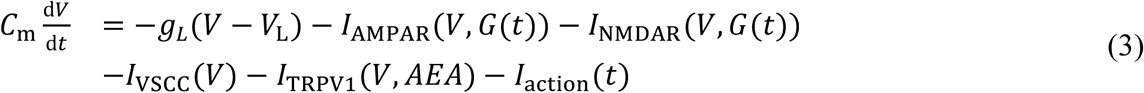

where *V* is the postsynaptic membrane potential and *AEA* stands for anandamide concentration. The mathematical expressions used for the currents of eq. (3) are given in Cui et al. (2016). The dynamics of free cytosolic calcium C was computed according to:

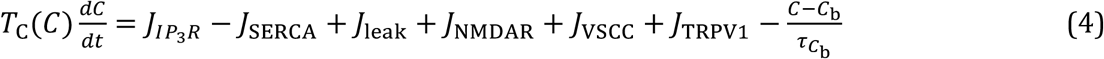

where *J*_NMDAR_, *J*_VSCC_ and *J*_TRPV1_ are the calcium fluxes from the respective plasma membrane channels (eq. 3). *T*_C_(*C*) is a time scaling factor accounting for the presence of endogenous calcium buffers. The terms 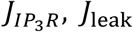 and *J*_*SERCA*_ describe calcium exchange flows between the cytosol and the endoplasmic reticulum (ER) via Inositol 1,4,5-trisphosphate (IP_3_)-Receptor channels (IP_3_R), passive ER-to-cytosol leak and Sarcoplasmic/Endoplasmic Reticulum Ca2^+^- ATPases (SERCA).

IP_3_ and diacylglycerol (DAG) are central signaling molecules in the model because they activate IP_3_R, thus increasing cytosolic Ca and allow the formation of 2-arachidonoylglycerol (2-AG) by calcium-activated DAG Lipase-α. Their dynamics is modeled as:

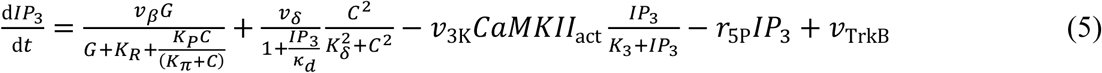

and

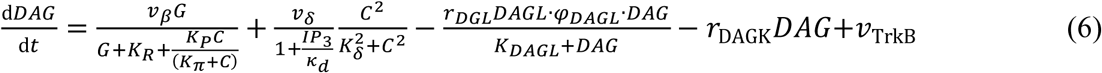

In both eq. (5) and (6), the first two terms of the right-hand size member correspond to IP_3_ and DAG synthesis via mGluR-activated PLCβ and Ca-activated (agonist independent) PLCδ. The third and fourth terms of the right-hand size member account for IP_3_ consumption by active CaMKII (*CaMKII*_act_) and Inositol 5-Phosphatase (eq. 5) or DAG consumption by DAG Lipase and DAG kinase (eq. 6).

The last term of eq. (5) and (6), i.e. *υ*_TrkB_ constitutes the main modification of the model implemented in the present study (i.e. *υ*_TrkB_ =0 in Cui et al. 2016). This term represents a constant production of IP_3_ and DAG by TrkB and accounts, using the simplest formulation, for TrkB-triggered activation of PLCγ. Therefore, our model includes the effect of BDNF and TrkB on IP_3_ and DAG production by PLCγ, thus disregarding any other potential effects on corticostriatal plasticity (mitogen-activated protein kinase, MAPK, or phosphatidylinositol 3-kinase, PI3K, pathways).

Our model also accounts for the biochemical pathways leading to the production of the endocannabinoids 2-AG and AEA, and their subsequent activation of cannabinoid receptors type-1, CB_1_R (see Fig. 5A). In agreement with experimental observation (Cui et al, 2015), CB_1_R activation in the model sets the synaptic weight *W* in a biphasic way: moderate amount CB_1_R activation decrease *W* while large amounts of CB_1_R activation increase it. This principle was modeled using a phenomenological expression:

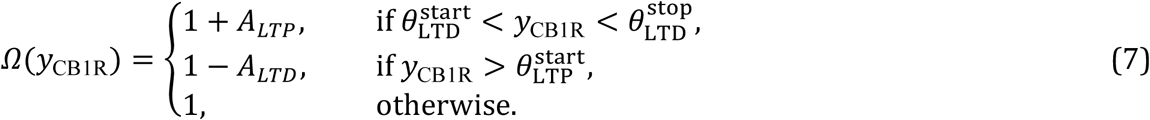

where 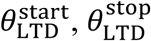 and 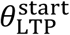 are plasticity thresholds and *y*_CB1R_ is proportional to the fraction of activated CB1R. Finally, the function *Ω*(*y*_CB1R_) sets the synaptic weight *W* according to:

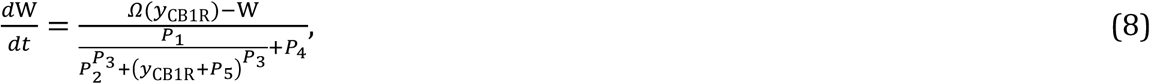

#### Parameters and numerics

We set the values of almost all the model parameters to their values in Cui et al (2016), except for 6 parameters related to IP_3_ and DAG dynamics, that we altered to account for the addition of *υ*_TrkB_ in eq. (5) and (6) above: (*υ*_TrkB_, *κ*_*d*_, *K*_δ_, *K*_*R*_, υ_δ_, *CaM*_tot_), representing respectively the rate of production of IP_3_ by TrkB, PLCδ product inhibition, PLCδ Ca-activation, glutamate affinity to mGluR, PLCδ maximal rate and the total concentration of calmodulin. Estimation of those 6 parameters was achieved using Differential Evolution (pagmo2 python library) to minimize the least-square distance between model outputs and experimental measurements of *W*_*total*_ obtained with various values of the STDP protocol parameters (*N*_pairings_, Δ*t*_STDP_, *S*_TrkB_). Here *S*_TrkB_={DHF, ANA12, Ctrl} specifies the pharmacology used against TrkB, i.e. addition of DHF/TrkB agonist, addition of ANA12 or K252a/TrkB antagonist or no addition. To reflect the importance of eCB-tLTP in our study, the weights of the corresponding data points (i.e. *N*_pairings_=25, Δ*t*_STDP_=-15 ms) were fivefold that of the other data points. The estimated parameter values were *κ*_*d*_=0.0623 µM, *K*_γ_=0.3514 µM, *K*_*R*_=4.9999 µM, υ_δ_=0.0209 µM/s, *CaM*_tot_=0.0714 µM and *υ*_TrkB_ =0.010 µM/s (*S*_TrkB_=Ctrl), 0.000 µM/s (*S*_TrkB_=ANA12) or 0.015 µM/s (*S*_TrkB_=DHF).

Numerical integration was performed as detailed in Cui et al (2016), i.e. using the ODEPACK LSODA solver compiled from fortran77 for python with f2py. Initial conditions were set to the steady-state of each variable in the absence of stimulation.

## RESULTS

We investigated the involvement of TrkB activation in bidirectional eCB-plasticity induced by three distinct Hebbian activity patterns: (*i*) the eCB-LTD induced by a low frequency stimulation (LFS) protocol (600 cortical stimulations at 1 Hz) (Fino et al., 2005; Puente et al., 2011), (*ii*) the eCB-tLTD induced with 100 STDP pairings (Shen et al., 2008; Fino et al., 2010; Paillé et al., 2013; Cui et al., 2015) and (*iii*) the eCB-tLTP induced by low numbers of STDP pairings (*i.e.* ~10) (Cui et al., 2015; Cui et al., 2016; Cui et al., 2018; Xu et al., 2018). To do so, we performed whole-cell recordings from MSNs of the dorsolateral striatum in horizontal brain slices (in which stimulations were performed in the layer 5 of the somatosensory cortex) at postnatal days P_25-35_ (Fig. 1A, 1B). Indeed, we have previously characterized these three forms of eCB-plasticity (LFS-LTD, eCB-tLTD and eCB-tLTP) at the level of the dorsolateral striatum in the very same preparation (Fino et al., 2005; Cui et al., 2015; Xu et al., 2018).

**Figure 1:**
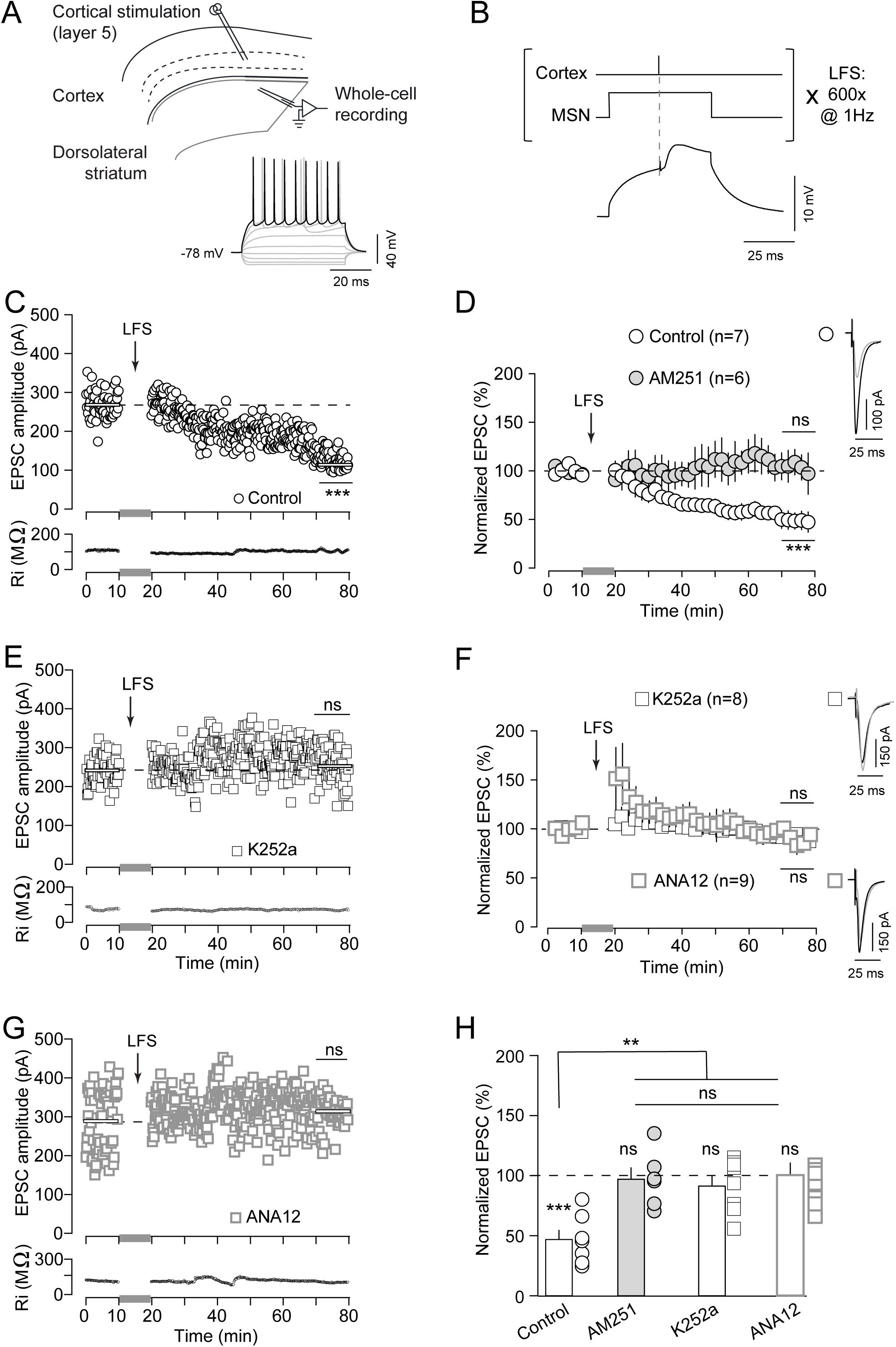
eCB-LTD induced with LFS relies on TrkB activation. (**A**) Scheme of the recording and stimulating sites in cortico-striatal horizontal slices and LFS protocol. (**B**) LFS protocol: 600 cortical stimulations at 1 Hz were paired with concomitant postsynaptic subthreshold depolarizations. (**C**-**D**) Corticostriatal eCB-mediated LTD induced by LFS. (**C**) Example of LTD induced by LFS the mean baseline EPSC amplitude was 268±4pA before LFS and decreased by 60% to 106±1pA one hour after LFS protocol (p=0.0001). Bottom, time course of Ri (baseline: 109±0.4MΩ and 50-60 min after pairings: 106±1MΩ; change of 3.6%) for this cell. (**D**) Averaged time-courses of LTD induced by LFS (7/7 cells showed LTD); this LFS-LTD was dependent on CB_1_R activation, because AM251 (3 µM) prevented tLTD (2/6 cells showed LTD). (**E**-**G**) Corticostriatal LFS-LTD relies on TrkB activation. (**E**) Example of the lack of LTD after LFS in presence of K252a (200 nM) (240±4pA before LFS and a lack of plasticity was observed one hour after LFS: 254±6pA, *p*=0.0561). Bottom, time course of Ri (baseline: 75±1MΩ and 50-60 min after pairings: 74±0.2MΩ; change of 1.4%). (**F**) Averaged time-courses showing the lack of LTD after LFS in presence of K252a (3/8 cells showed LTD) or with ANA12 (3/9 cells showed LTD). (**G**) Example of the lack of LTD after LFS in presence of ANA12 (10 µM) (286±11pA before LFS and 310±5pA one hour after LFS, *p*=0.0735). Bottom, time course of Ri (baseline: 98±1MΩ and 50-60 min after pairings: 93±1MΩ; change of 5%). (**H**) Graph bars of the averaged responses for all control, AM251, K252a and ANA12 conditions (One-way ANOVA, F_(3,26)_ = 7.784; *p*=0.0007). Insets (in **D** and **F**) correspond to the average EPSC amplitude during baseline (black trace) and during the last 10 min of recordings after LFS (grey trace). Error bars represent the SEM. ***: *p*<0.001; ns: not significant.

### TrkB activation is necessary for eCB-LTD induced by LFS

The LFS protocol consisted in 600 cortical stimuli at 1 Hz with concomitant depolarization (50 ms) of the recorded MSNs (Fig. 1B) and induced reliable LTD (LFS-LTD), in line with previous studies (Fino et al., 2005; Puente et al., 2013). An example of LFS-LTD recorded during one hour after LFS is shown in Figure 1C; Input resistance (Ri) remained stable over this period. Overall, LFS induced LTD (mean value of EPSC amplitude recorded 50 min after LFS protocol induction: 47±8%, *n*=7, *p*=0.0005). This striatal LFS-LTD was CB_1_R-mediated since prevented by AM251 (3μM), a specific CB_1_R inhibitor (97±9%, n=6, *p*=0.7729) (Fig. 1D).

We then investigated whether TrkB activation was required for the induction of the LFS-LTD. Bath-applied K252a (200 nM), a selective inhibitor of the tyrosine kinase activity of the Trk family, prevented LFS-LTD. Figure 1E shows an example of the absence of synaptic plasticity with K252a. In summary, K252a prevented the induction of LFS-LTD (91±7%, *p*=0.2101, n=8) (Fig. 1F and 1H). To confirm this finding, we then used ANA12 (10 µM), a competitive antagonist of TrkB, structurally distinct from K252a. In the example shown in the Figure 1G, LFS with bath-applied ANA12 failed to induce LTD. In summary, we observed that ANA12 prevented the induction of LFS-LTD (100±10%, *p*=0.9725, n=9) (Fig. 1F and 1H).

Altogether our results demonstrate that TrkB activation is necessary for LFS-eCB-LTD expression (Fig. 1H; One-way ANOVA *p*=0.0007).

### STDP-induced eCB-tLTD requires TrkB activation

We next tested whether TrkB activation was necessary for another form of eCB-LTD, a spike-timing-dependent LTD (tLTD). This plasticity is induced with a STDP protocol consisting in pairing the pre- and postsynaptic stimulations separated by a fixed temporal interval, Δt_STDP_, repeated here 100 times at 1 Hz (with Δt_STDP_~15ms) (Fig. 2A). We have previously shown that GABA operates as a Hebbian/anti-Hebbian switch at corticostriatal synapses (Paillé et al., 2013; Valtcheva et al., 2017) and corticostriatal STDP polarity depends on the presence (*in vitro* Hebbian STDP; Pawlak and Kerr, 2008; Shen et al., 2008) or absence (*in vitro* anti-Hebbian STDP; Fino et al., 2005; Fino et al., 2010; Cui et al., 2015; *in vivo* anti-Hebbian STDP; Schulz et al., 2010) of GABA_A_ receptor antagonists. As exemplified in Supplementary Figure 1A, we observed that 100 pre-post pairings triggered corticostriatal tLTD. Overall, 100 pre-post pairings induced tLTD (68±7%, *p*=0.0010, *n*=10), which was CB_1_R-mediated since AM251 (3μM) fully prevented tLTD expression (103±7%, *p*=0.5917, *n*=5) (Fig. 2B), in agreement with previous results (Pawlak and Kerr, 2008; Shen et al., 2008; Fino et al., 2010; Cui et al., 2015).

**Figure 2:**
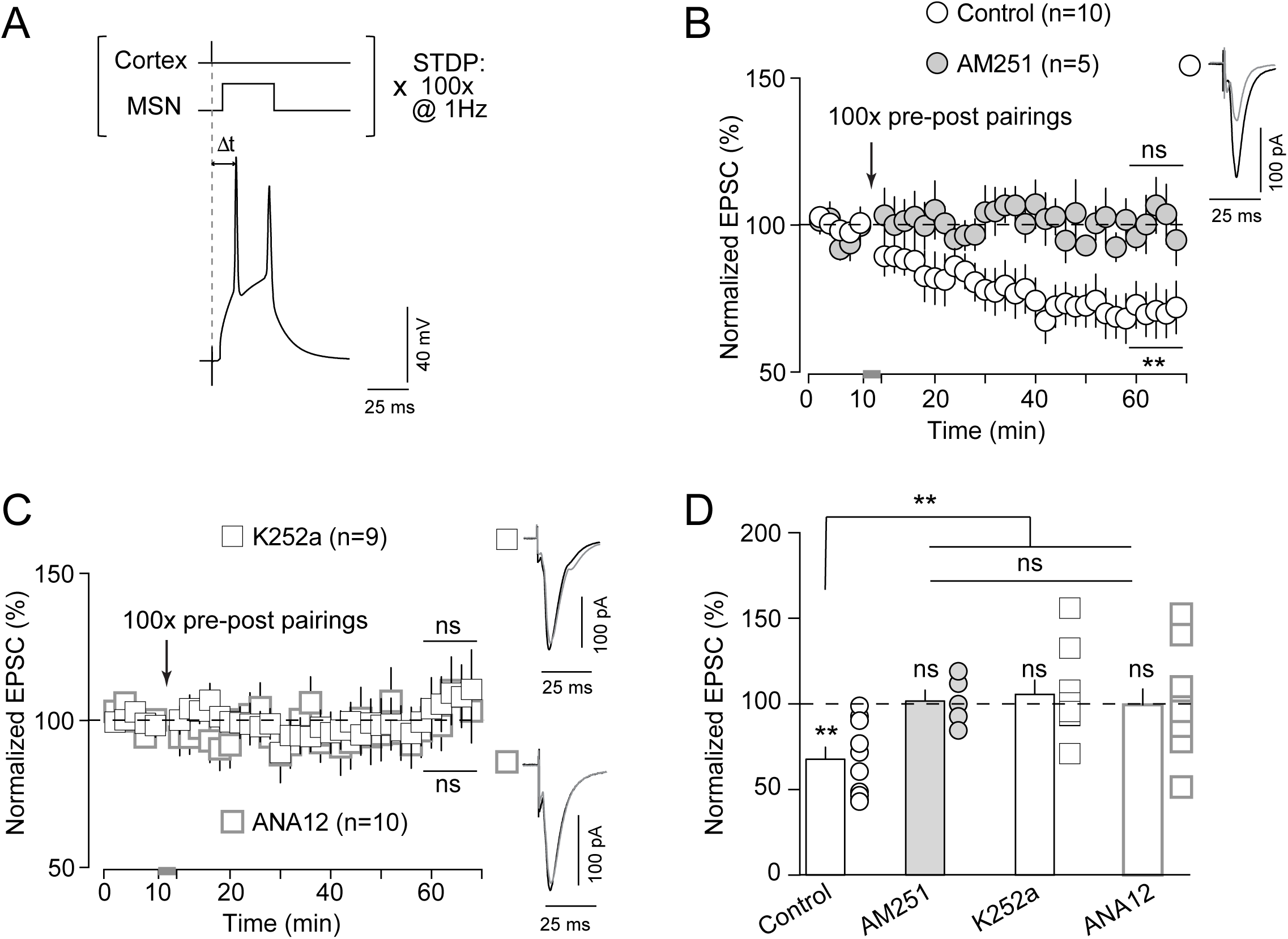
eCB-tLTD relies on TrkB activation. (**A**) Scheme of the recording and stimulating sites in corticostriatal horizontal slices and the STDP protocol: a single cortical stimulation was paired with a single spike evoked by a depolarizing current step in the recorded striatal MSN, in a pre-post sequence (0<Δt_STDP_<+20 ms); this pairing was repeated 100 times at 1 Hz. (**B**) Corticostriatal eCB-mediated tLTD. Averaged time-courses of tLTD induced by 100 pre-post pairings (8/10 cells displayed tLTD); this tLTD was mediated by CB_1_R, because tLTD was prevented by the application of AM251 (3 µM) (1/5 cells showed tLTD). (**C**) Corticostriatal eCB-tLTD relies on TrkB activation. Averaged time-courses showing the lack of plasticity observed with K252a (1/9 showed tLTD) and with ANA12 (3/10 showed LTD). (**D**) Graph bars of the averaged responses for all control, AM251, K252a and ANA12 conditions (One-way ANOVA, F_(3,30)_ = 4.84; *p*=0.0073). Insets (**B** and **C**) correspond to the average EPSC amplitude during baseline (black trace) and during the last 10 min of recording after STDP pairings (grey trace). Error bars represent the SEM. **: *p*<0.01; ns: not significant.

We next investigated whether the activation of TrkB was required for the induction of the eCB-tLTD. K252a (200 nM) application prevented eCB-tLTD expression, as exemplified in the Supplementary Figure 1B. Overall, K252a prevented eCB-tLTD (105±8%, *p*=0.5847, n=9) (Fig. 2C and 2D). Similarly, ANA12 (10 uM) prevented eCB-tLTD, as exemplified in the Supplementary Figure 1C. In summary, ANA12 abolished eCB-tLTD expression (101± 9%, *p*=0.9699, n=10) (Fig. 2C and 2D). Altogether this set of results reveals that eCB-tLTD is dependent on the activation of TrkB (Fig. 2D; One-way ANOVA *p*=0.0073).

### STDP-induced eCB-tLTP requires TrkB activation

We have previously reported that low numbers of STDP pairings (~10) induce an eCB-tLTP, a form of plasticity dependent on the activation of CB_1_R at corticostriatal synapses in the dorsolateral striatum (Cui et al., 2015; Cui et al., 2016; Cui et al., 2018; Xu et al., 2018). We thus tested whether the involvement of TrkB was crucial not only in eCB-mediated depression (LFS-eCB-LTD and eCB-tLTD) but also in eCB-tLTP. 10 post-pre pairings at 1 Hz (Fig. 3A) induced tLTP as exemplified in Supplementary Figure 1D. Overall, 10 post-pre STDP pairings induced tLTP (153±21%, *p*=0.0021, n=9), which was prevented by AM251 (3μM) (77±9%, p=0.0508, n=6) (Fig. 3B).

**Figure 3:**
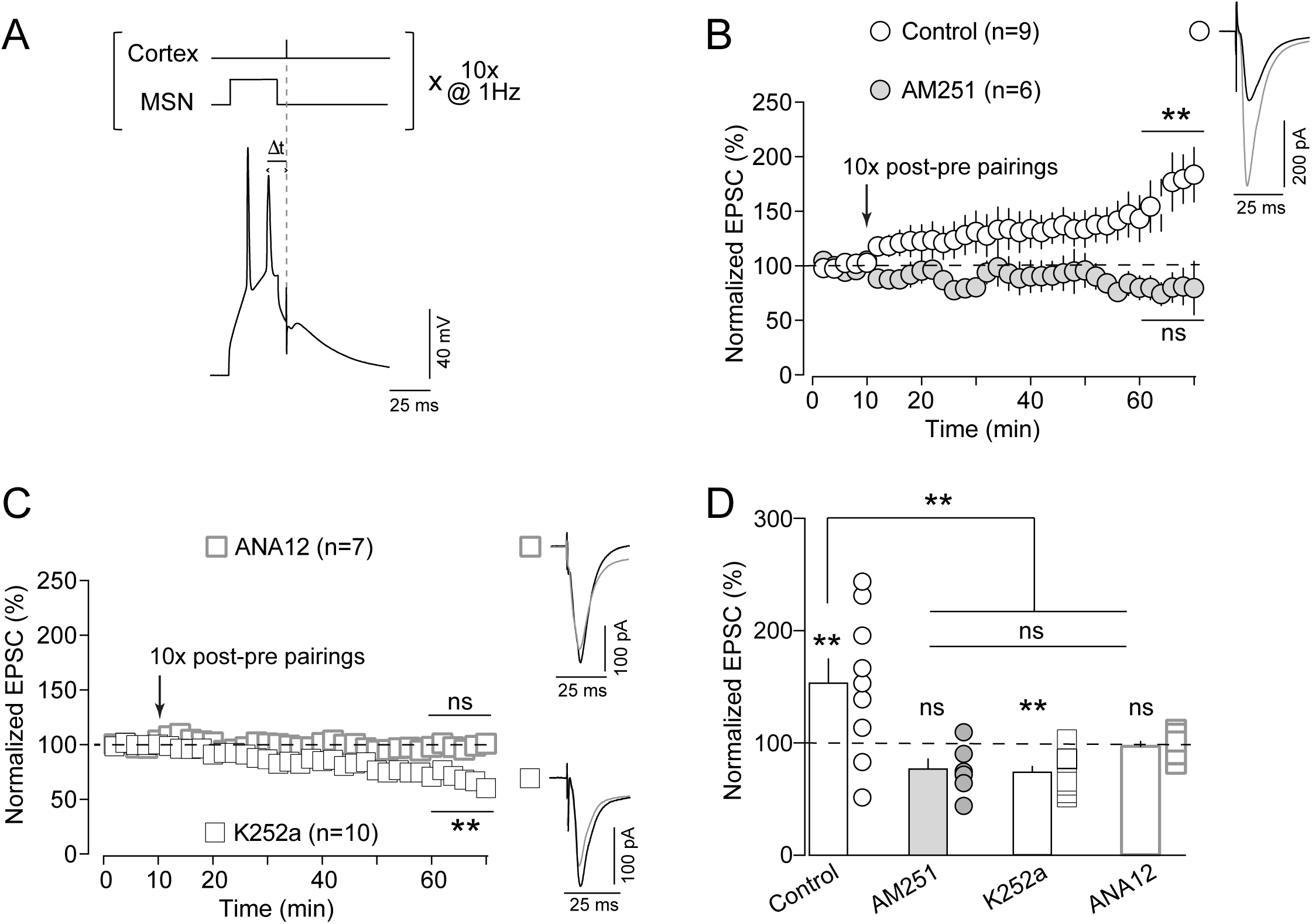
eCB-tLTP relies on TrkB activation. (**A**) Scheme of the recording and stimulating sites in cortico-striatal horizontal slices and the STDP protocol: a single cortical stimulation was paired with a single spike evoked by a depolarizing current step in the recorded striatal MSN, in a post-pre sequence (−20<Δt_STDP_<0 ms); this pairing was repeated 10 times at 1 Hz. (**B**) Corticostriatal eCB-mediated tLTP. Averaged time-courses of tLTP induced by 10 post-pre pairings (6/9 cells showed tLTP); this tLTP was mediated by CB_1_R, because tLTP was prevented by the application of AM251 (3 µM) (0/6 showed tLTP). (**C**) Corticostriatal eCB-tLTP relies on TrkB activation. Averaged time-courses showing the lack of plasticity observed with K252a (0/10 showed tLTP, and 8/10 showed tLTD) or with ANA12 (0/7 showed tLTP). (**D**) Graph bars of the averaged responses for all control, AM251, K252a and ANA12 conditions (One-way ANOVA, F_(3,28)_ = 8.60; *p*=0.0003). Insets (**B** and **C**) correspond to the average EPSC amplitude during baseline (black trace) and during the last 10 min of recording after STDP pairings (grey trace). Error bars represent the SEM. ***: *p*<0.001; ns: not significant.

Blockade of TrkB by K252a (200 nM) prevented eCB-tLTP induction, as exemplified in Supplementary Figure 1E. Overall, K252 prevented the induction of eCB-tLTP and even induced tLTD (74±5%, *p*=0.0006, n=10) (Fig. 3F; see also Discussion). Using ANA12, we also observed that 10 post-pre pairings did not induce tLTP, as exemplified in the Supplementary Figure 1F. In summary, ANA12 prevented eCB-tLTP (97±4%, *p*=0.4827, n=7) (Fig. 3C and 3D). Altogether these results show that BDNF/TrkB signaling is necessary for corticostriatal eCB-tLTP (Fig. 3D; One-way ANOVA p=0.0003).

In conclusion, bidirectional eCB-mediated plasticity, i.e. eCB-LTD (induced by LFS or STDP induction protocol) as well as eCB-tLTP (induced with low numbers of STDP pairings), is dependent on TrkB activation.

### Activation of TrkB prolongs calcium transients

eCBs (2-AG and AEA) are synthesized and released on-demand (Alger and Kim, 2011). Enzymes involved in eCB biosynthesis exhibit a tight dependence on cytosolic calcium and consequently eCB-mediated plasticity strongly relies on calcium dynamics (Chevaleyre et al., 2006, Heifets and Castillo, 2015). It exists a link between TrkB intracellular signaling pathways and calcium dynamics via the activation of the PLC signaling pathway, and more precisely PLCγ (Hashimotodani et al., 2005).

Here, we examined whether TrkB activation induced a change in cytosolic Ca^2+^-evoked events, which could account for its involvement in eCB-mediated plasticity. We monitored Ca^2+^-evoked events in dendritic spines and adjacent shafts using two-photon microscopy in line-scan mode (Fig. 4A1 and A2) using ratiometric indicators Fluo-4F (250µM) and Alexa Fluor 594 (50µM) (Fig. A3). We examined calcium transients under TrkB activation (using the specific TrkB agonist DHF, 10µM) while eliciting either two back-propagating action potentials (bAPs) without presynaptic paired stimulation (Fig. 4B), or pairing one corticostriatal EPSP with two bAPs for Δ*t*~+15ms (Fig. 4C) or for Δ*t*~-15ms (Fig. 4D).

**Figure 4:**
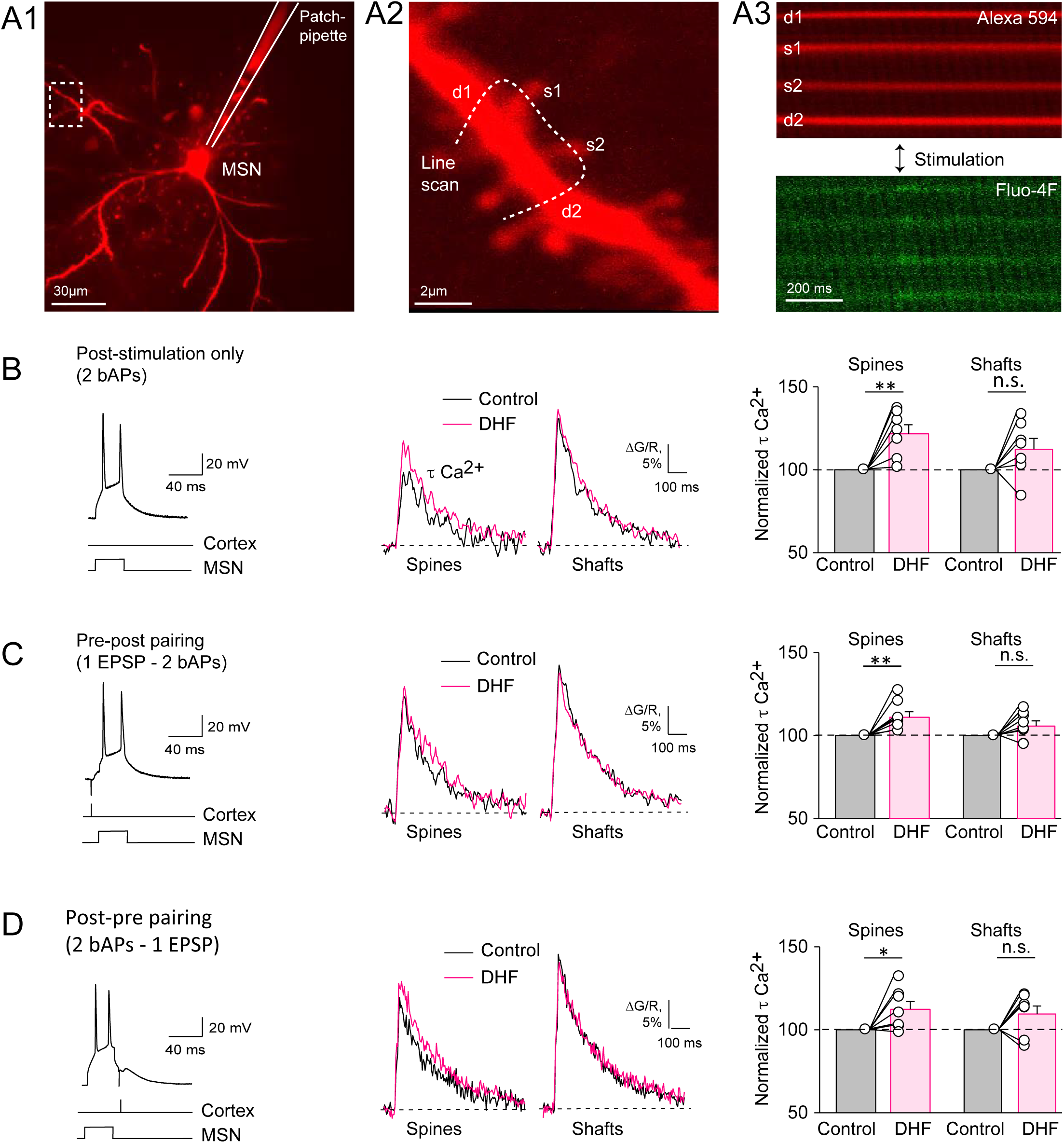
Activation of TrkB prolongs calcium transients in dendritic spines. (**A**) (**A1**) experimental set-up showing the combination of whole-cell patch-clamp of a single MSN (see the patch-clamp pipette underlined in white) with two-photon imaging; the dashed white square indicated the imaged area). The scanning areas had been selected on 100-150 µm distance from soma. Scale bar: 30 µm. (**A2**) Line-scanning two-photon microscopy of Ca^2+^ transients in dendritic spines and adjacent shafts in MSNs filled with (**A3**) ratiometric indicators Fluo-4F (250 µM) and Alexa Fluor 594 (50 µM). A2: scale bar: 2 µm; A3: scale bar: 2 ms. (**B**-**D**) DHF increased Ca^2+^ transients in dendritic spines, but not in shafts, triggered by (**B**) two bAPs (post-stimulation only), (**C**) paired stimulation, consisting in single evoked corticostriatal EPSP paired with two bAPs with Δ*t* ~+15-20 ms, or (**D**) paired stimulation, consisting two bAPs paired with single evoked corticostriatal EPSP with Δ*t* ~-15-20 ms. Error bars represent the SEM. *: *p*<0.05; **: *p*<0.01; ns: not significant.

First, during unpaired postsynaptic stimulation (bAPs only) we found that application of DHF increased the decay τ constant of Ca^2+^ response (τCa^2+^) to 124±5% of control (n=7, *p*=0.0060) in spines whereas no changes of τCa^2+^ was observed in adjacent dendritic shafts (112±6% of control, n=7, *p*=0.0943) (Fig.4B). We next pre-treated with ANA12 (10 µM) and then applied DHF to ensure that DHF-induced effects on τCa^2+^ were TrkB-mediated (Supplementary Fig. 2A). In both spines and shafts, we observed that τCa^2+^ remained unaffected after ANA12 application (spines: 103±3%, n=8, *p*=0.3099; shafts: 103±1%, *p*=0.0516) or after ANA12+DHF co-application (spines: 102±4% of control, *p*=0.8776; shafts: 108±2%, *p*=0.0704). Since DHF and ANA12 were dissolved in DMSO (0.02-0.04% final concentration), we tested the effects of DMSO alone to ensure the specificity of the DHF and ANA12 effects (Supplementary Fig. 2B). In spines and shafts we found no significant difference in τCa^2+^ between control and DMSO (spines: 100±3% n=7, *p*=0.9882; shafts: 103±4%, *p*=0.4444).

Second, we applied paired stimulations consisted of cortical stimulation followed by postsynaptic bAPs (Δ*t*~+15ms), i.e. pre-post pairings as used for eCB-tLTD induction (Fig. 4C). Under DHF application, pre-post pairings induced an increase in τCa^2+^ in dendritic spines to 111±3% when compared to control (n=8, *p*=0.0085) (Fig. 4C). In dendritic shafts, we did not observe significant changes in τCa^2+^ (106±3%, n=8, *p*=0.0847) (Fig. 4C). In both dendritic spines and shafts, we found that τCa^2+^ were not modified by ANA12 treatment (spines: 104±3%, n=6, *p*=0.1975; shafts: 98±2%, p=0.4030) or when ANA12 and DHF were co-applied (spines: 105±7%, *p*=0.8731; shafts: 99±6%, *p*=0.7712) (Supplementary Fig. 4C).

Third, we tested whether post-pre pairings yielded similar results than with bAPs only or with pre-post STDP pairings (Fig. 4D). To do so, we applied pairings consisting of postsynaptic bAPs followed by cortical stimulation (Δ*t*~-15ms). Under DHF application, post-pre pairings induced an increase in τCa^2+^ in dendritic spines to 112±5% compared to control (n=7, *p*=0.0391) (Fig. 4D). In dendritic shafts, we did not observe significant changes in τCa^2+^ (109±5%, n=7, *p*=0.0941) (Fig. 4D).

We did not observe a significant difference in the amplitude of Ca^2+^-evoked events along recording time and between DHF, ANA12 and DMSO applications compared to control during postsynaptic stimulation only (2bAPs) (Supplementary Figure 3A) or during presynaptic stimulation (EPSP) paired with postsynaptic stimulation (2bAPs) (Supplementary Figure 3B). Altogether these results indicate that TrkB activation modifies the dynamics of the Ca^2+^ transients (triggered by bAPs only, pre-post or post-pre STDP pairings), and promotes longer Ca^2+^ events with higher quantity of intracellular calcium. This increased calcium signaling upon TrkB activation can be viewed as a necessary boost to favor the synthesis and release of eCBs.

### Predicting the effect of TrkB activation on eCB-STDP with a mathematical model

We questioned how TrkB activation could participate to eCB-tLTD and eCB-tLTP, depending on the activity pattern of either side of the synapse. To address this question, we built a mathematical model of the molecular mechanisms of corticostriatal synaptic plasticity (Fig. 5A). Our model is based on the NMDAR-and CB_1_R-signaling pathways involved in corticostriatal STDP (Pawlak and Kerr, 2008; Shen et al., 2008; Fino et al., 2010; Paillé et al., 2013; Cui et al., 2015) and actually extends to TrkB signaling the mathematical model we validated in previous studies (Cui et al., 2016; Cui et al., 2018; Xu et al., 2018). It expresses the kinetics of the enzymes and binding reactions implicated, including the effects of AMPAR, NMDAR, VSCC and TRPV1, on cytosolic Ca^2+^, IP_3_-controled Ca^2+^-induced Ca^2+^ release from the endoplasmic reticulum, the synthesis of endocannabinoids (2-AG and AEA) via DAG lipase α and FAAH pathways, respectively, and the retrograde activation of presynaptic CB_1_R (Fig. 5A). In agreement with experimental evidence (Cui et al., 2015; Cui et al., 2016; Cui et al., 2018; Xu et al., 2018), CB_1_R activation sets the value of the synaptic weight *W* in the model according to a biphasic mechanism: moderate amounts of CB_1_R activation yield LTD whereas high level of CB_1_R activation yields LTP. Note that such a concentration-dependent biphasic control of the signaling pathway downstream of CB_1_R has consistently been evidenced *in vitro* (Glass et al., 1997, Jarrahian et al., 2004, Kearn et al., 2005, Eldeeb et al., 2016).

Our model contains three isoforms of phospholipase-C (PLCβ, δ and γ) in the postsynaptic neuron: PLCβ is activated by binding of presynaptically-released glutamate to mGluR, a G_q/11_-coupled GPCR; PLCδ is agonist-independent but tightly controlled by cytosolic Ca^2+^; and PLCγ is activated by TrkB (Fig. 5A). In spite of those distinct properties, the three PLC catalyze the same reaction: the synthesis of IP_3_ and DAG from phosphatidylinositol biphosphate (PIP_2_). In turn, IP_3_ activates IP_3_Rs channels at the membrane of the endoplasmic reticulum, which boost the calcium influx in the cytosol. Model predictions show that the transient Ca^2+^response to glutamate release upon presynaptic stimulation is wider (and slightly larger) when TrkB is activated when compared to control conditions (Fig. 5B), whereas it remains unchanged upon TrkB inhibition (Fig. 5B). This behavior matches the calcium two-photon imaging experiments reported in the Figure 4 with the use of TrkB agonist and antagonist, DHF and ANA12, respectively.

In control conditions (Fig. 5C1), the model reproduces the characteristics of eCB-STDP expression: eCB-tLTD starts to be expressed when at least 25 pre-post stimulations are applied (with Δt ~+20ms) and progressively reinforces when *N*_pairings_ increases. eCB-tLTP is observed for low numbers of post-pre pairings (*N*_pairings_ ∈ [2,20], Δt ~-15ms) and disappears after 20 pairings.

In the model, the activation of TrkB (Fig. 5C2) has two main effects following STDP induction protocols: (*i*) the eCB-tLTP region (red region in Fig. 5C2) expands considerably in terms of Δt and *N*_pairings_ and (*ii*) eCB-tLTD is more pronounced (i.e. the blue regions of tLTD in Fig. 5C2 are darker than those in Fig. 5C1). On the contrary, the inhibition of TrkB precludes the induction of eCB-tLTP (note the lack of red region in Fig. 5C3) and considerably weakens eCB-tLTD (Fig. 5C3). Comparison of model outputs with experimental data can be carried out for the (*N*_pairings_, Δt) values that were tested experimentally.

In the model, the effects of TrkB activation and inhibition can be understood because of the threshold nature of eCB-STDP expression: eCB-tLTP is expressed when large amounts of 2-AG are produced and CB_1_R is activated above the tLTP threshold (*θ*_LTP_^start^, Fig. 5D). In control conditions, CB_1_R activation overcomes *θ*_LTP_^start^ only for *N*_pairings_>6 and up to *N*_pairings_<20 (for larger *N*_pairings_, CB_1_R desensitization strongly reduces CB_1_R activation), therefore eCB-tLTP is expressed for 6<*N*_pairings_<20 (Fig. 5C1). Part of the amount of 2-AG needed for CB_1_R activation is contributed by TrkB via IP_3_ production by PLCγ. For example, when TrkB is completely inhibited, the amount of IP_3_ produced at each pairing is lower compared to control conditions, which in turn decreases 2-AG production and consequent activation of CB_1_R (Fig. 5D, *green* line). Under TrkB inhibition, 2-AG production may not be sufficient to overcome the LTP threshold, thus explaining the disappearance of eCB-tLTP observed with ANA12 (Fig. 3C and 3D). With complete inhibition of TrkB, the decrease of CB_1_R activation can even be large enough that CB_1_R activation enters the LTD range (Fig. 5D, *green* line). In this case, one expects a shift of plasticity, with eCB-tLTP shifting to eCB-tLTD, in agreement with the K252a experiments (Fig. 3D). The same reasoning explains that eCB-tLTD is considerably weakened under TrkB inhibition, as well as the effects of DHF.

Figure 5E1 shows the close agreement between model predictions and synaptic efficacy changes in control conditions and with the inhibition of TrkB (“TrkB inhibited” conditions). In particular, under Trkb inhibition, the model faithfully reproduces the disappearance of eCB-tLTD at 100 pre-post pairings (shown experimentally in Figure 2) and that of eCB-tLTP at 10 post-pre pairings (shown experimentally in Figure 3).

The above results (Fig. 5C and E1) validate the model. We then used the model to derive experimentally-testable predictions about the effect of TrkB agonists (such as DHF). Figure 5E shows model prediction for the change of synaptic weight *W* with *N*_pairings_ using STDP protocols yielding eCB-tLTP (upper curve, Δt=-20 ms) and eCB-tLTD (lower curve, Δt=+17ms). The model predicts that DHF should increase the amplitude of eCB-tLTP but also the range of *N*_pairings_ values where it can be induced. The amplitude of eCB-tLTD should be increased, at least for *N*_pairings_>25, with DHF (Fig. 5E2). Note that, in the model, the effects of TrkB are restricted to the activation of PLCγ and resulting changes in the production of IP_3_ and DAG. The model does not integrate the other pathways of TrkB signaling (e.g. MAPK or PI3K). Therefore, experimental validation of the above predictions would strongly suggest that the effects of TrkB on eCB-STDP are mainly underlain by PLCγ.

### Activation of TrkB promotes eCB-plasticity

We further investigated the causal role of TrkB activation in bidirectional eCB-plasticity and tested the model predictions (Fig. 5E). We tested two scenarios in which the boost in eCB synthesis operated by TrkB activation (with DHF, 10µM) should expand eCB-STDP domains for (*i*) eCB-tLTD (for N_pairings_<50) and (*ii*) eCB-tLTP (for N_pairings_>20). For this purpose, we chose two STDP protocols involving 25 pre-post and 25 post-pre pairings, for which eCB-tLTD and eCB-tLTP, respectively, are not observed in control conditions (Cui et al., 2015; Cui et al., 2016).

First, we tested the effect of the activation of TrkB by DHF for 25 pre-post pairings. In control conditions, an absence of plasticity was observed following 25 pre-post pairings (99±3%, *p*=0.9728, n=6) (Fig. 6A). This result is in line with our previous reports showing that eCB-tLTD progressively disappeared when N_pairings_<50 (Cui et al., 2015; Cui et al., 2016). Following DHF application, a tLTD was observed for 25 pre-post STDP pairings (61±5%, *p*=0.0001, n=7) (Fig. 6A and 6C). This tLTD was CB_1_R-mediated since its expression was prevented by AM251 (3µM) (103±6%, *p*=0.6287, n=6) (Fig. 6B and 6C). Besides the PLCγ pathway, TrkB activation also leads to the recruitment of the MAPK and PI3K pathways. Thus, we next tested whether this DHF-induced eCBtLTD could still be induced following pharmacological blockade of the MAPK and PI3K pathways. For this purpose, specific inhibitors of MAP kinase kinases (MEK1 and MEK2), U0126 (10µM), and of PI3K, LY294002 (10µM), were co-applied during the whole experiment including STDP pairings. Under inhibition of the MAPK and PI3K pathways, DHF was still able to induce tLTD for 25 pre-post pairings (58±5%, *p*=0.0007, n=6) (Fig. 6B and 6C).

**Figure 5:**
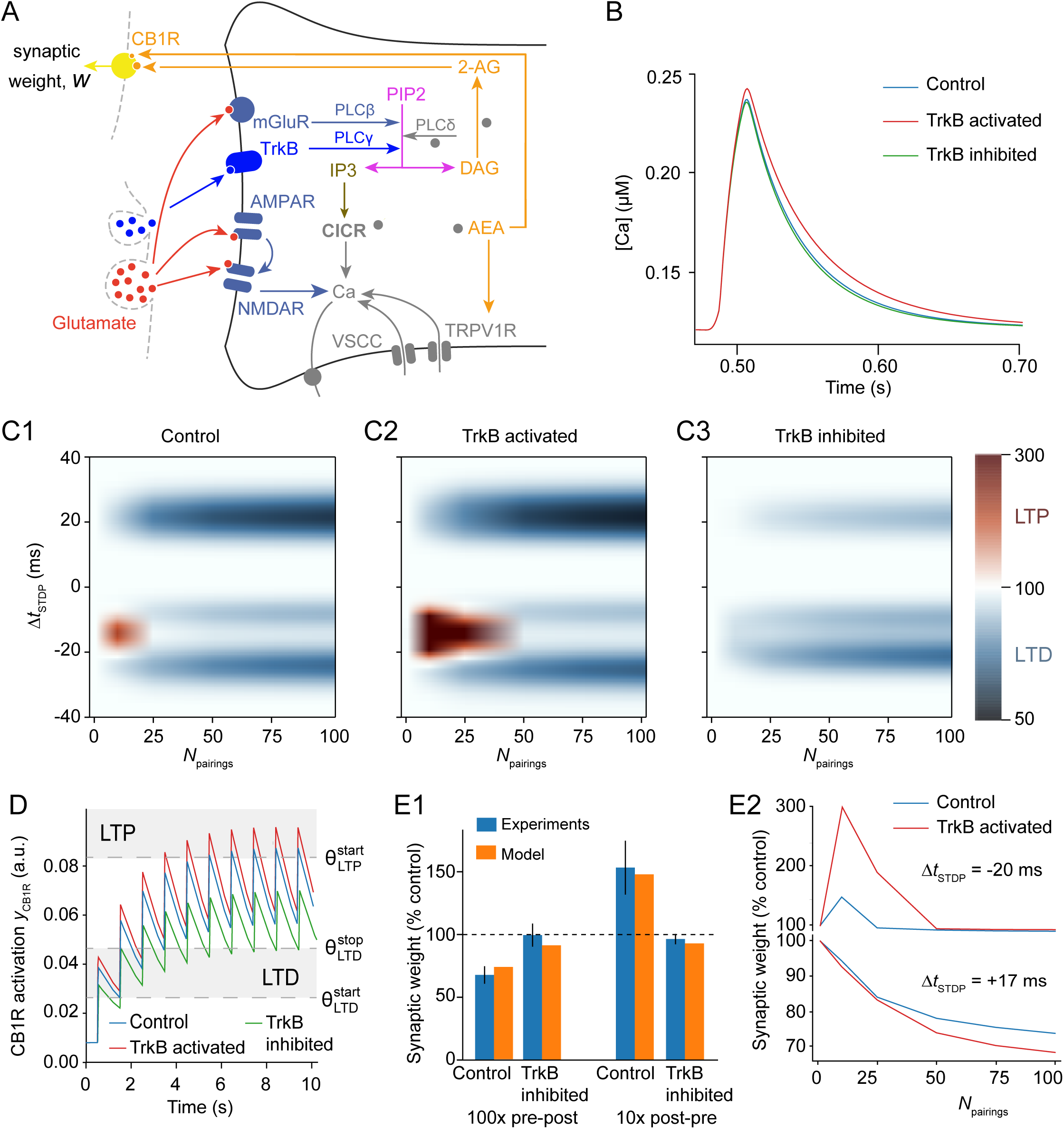
Effects of TrkB agonists in a mathematical model of the signaling pathways. (**A**) Scheme of the main signaling pathways accounted for in the mathematical model. The model expresses the kinetics of the corresponding enzymatic and binding reactions. To derive it, we used the mathematical model for corticostriatal STDP (Cui et al, 2016; Cui et al., 2018) and added the production of IP_3_ and DAG by TrkB-activated PLCγ. The full grey circles located the reactions that are controlled by cytosolic Ca^2+^. The synaptic weight W is defined by W_total=_W_post_×W_pre_. Abbreviations: CICR: Ca^2+^-induced Ca^2+^-release, mGluR: metabotropic glutamate receptor, AEA: anandamide; VSCC: Voltage-Sensitive Ca^2+^channels; TRPV1: Transient Receptor Potential cation channel subfamily V type 1. (**B**) Effect of the activation of TrkB and inhibition of TrkB on transients of cytosolic Ca^2+^ triggered by a single postsynaptic stimulation (to be compared with the experimental traces obtained in Fig. 4B). (**C**) Changes of the synaptic weight *W* in the model when the spike timing Δ*t*_STDP_ and the number of pairings of the STDP protocol are varied. *W* is analyzed in control conditions (**C1**), with TrkB activated (such as DHF-like conditions) (**C2**), and with TrkB inhibited (such as ANA12-or K252a-like conditions) (**C3**). The colorcode for those maps is given at the end of the line. (**D**) Time course of the model prediction for the activation of CB_1_Rs during the first 10 paired stimulations in control conditions (*blue*) and with activation of TrkB (*red*) or inhibition of TrkB (*green*). According to the model, tLTP is expressed when CB_1_R activation overcomes the threshold *θ*_LTP_^start^, whereas tLTD is expressed when CB_1_R activation is between *θ*_LTD_^start^ and *θ*_LTD_^stop^. Δ*t*_STDP_=-15 ms. (**E1**) Comparison of the effects of an inhibition of TrkB in the model (*blue*) with the experimental results (data with ANA12) (*orange*). (**E2**) Model predictions for the effects of the activation of TrkB on the changes of *W* with *N*_pairings_. In **E1** and **E2**, simulation for the model were performed for 100 pre-post pairings at Δ*t*_STDP_=+17 ms and for 10 post-pre pairings at Δ*t*_STDP_= −20 ms.

**Figure 6:**
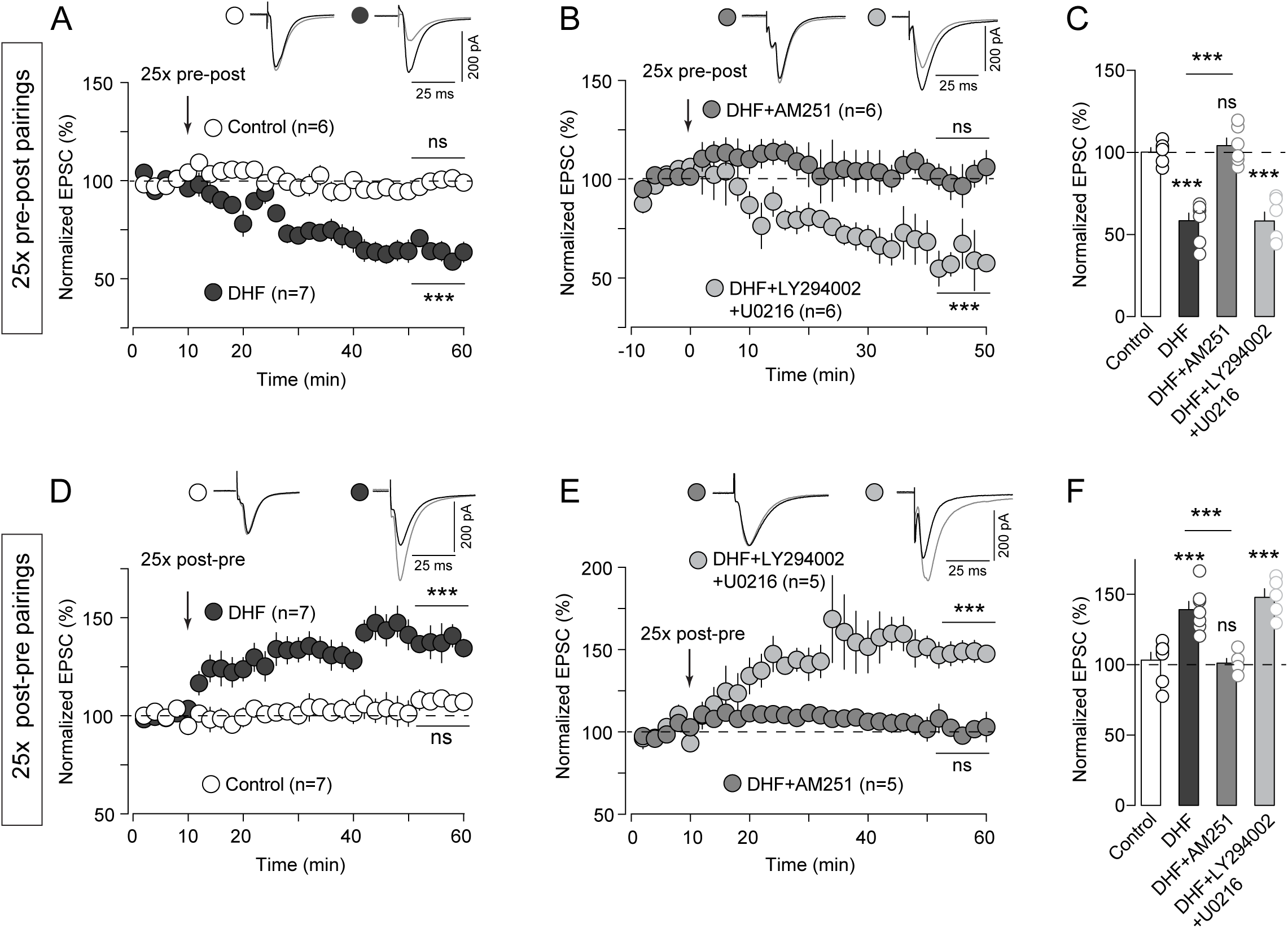
Activation of TrkB promotes eCB-plasticity. (**A**) Activation of TrkB with DHF (10µM) promotes eCB-tLTD for 25 pre-post pairings. Averaged time-courses showing the absence of plasticity (1/6 showed tLTD) in control conditions, and tLTD induced by 25 pre-post pairings with DHF bath-applied during STDP pairings (7/7 showed tLTD). (**B**) This tLTD was mediated by CB_1_R, because it was prevented by the application of AM251 (3µM) (0/6 showed tLTD). Inhibition of MAPK and PI3K pathways with U0126 (10µM) and LY294002 (10µM) did not prevent tLTD induced by 25 pre-post pairings with DHF (6/6 showed tLTD). (**C**) Summary bar graphs showing the eCB-tLTD induced by DHF together with 25 pre-post pairings and which is independent of the MAPK and PI3K pathways (One-way ANOVA, F_(3,21)_ = 32.44; *p*=0.0001). (**D**) DHF promotes eCB-tLTP for 25 post-pre pairings. Averaged time-courses showing the absence of plasticity (3/7 showed tLTP) observed after 25 pre-post pairings in control conditions, and tLTP induced by 25 post-pre pairings with DHF bath-applied during STDP pairings (7/7 showed tLTP). (**E**) this tLTP was mediated by CB_1_R, because it was prevented by AM251 (1/5 showed tLTP). Inhibition of MAPK and PI3K pathways with U0126 (10µM) and LY294002 (10µM) did not prevent tLTP induced by 25 post-pre pairings with DHF (5/5 showed tLTP). (**F**) Summary bar graphs showing the eCB-tLTP induced by DHF together with 25 post-pre pairings and which is independent of the MAPK and PI3K pathways (One-way ANOVA, F_(3,20)_ = 18.16; *p*=0.0001). Insets correspond to the average EPSC amplitude during baseline (black trace) and the last 10 min of recording after STDP pairings (grey trace). Error bars represent the SEM. ***: *p*<0.001; ns: not significant.

Second, we examined the effect of the activation of TrkB by DHF for the 25 post-pre STDP pairings. In control conditions, an absence of plasticity was observed after 25 post-pre pairings (101±5%, *p*=0.9238, n=7) (Fig. 6D). When we bath-applied DHF during the whole recording, no plasticity was induced after one hour (96±11%, *p*=0.7593, n=5) (Supplementary Fig. 4). We next restricted the application of DHF to the duration of the 25 STDP post-pre pairings. In these conditions, a reliable tLTP was observed (138±5%, *p*=0.0004, n=7) (Fig. 6D and 6F). This tLTP was CB_1_R-dependent since its expression was prevented by AM251 (3µM) (102±4%, *p*=0.6951, n=5) (Fig. 6E and 6F). This eCB-tLTP did not involve the MAPK and PI3K pathways. Indeed, 25 post-pre pairings, with concomitant activation of TrkB (DHF application), were still able to induce tLTP (148±6%, *p*=0.0014, n=5) even in presence of U0126 and LY294002 (Fig. 6E and 6F).

Therefore, eCB-plasticity appears to require the involvement of the PLCγ pathway following TrkB activation without the involvement of the MAPK and PI3K pathways.

In conclusion, these experiments confirm the model prediction that the activation of TrkB allows an enlargement of the domain of expression of eCB-tLTD and eCB–tLTP, indicating that TrkB efficiently gates but also shapes eCB-plasticity expression.

## DISCUSSION

In the present study, we investigated the mechanistic and functional interaction between TrkB and eCB signaling in eCB-mediated LTD and LTP at corticostriatal synapses. We found that activation of TrkB is a necessary upstream modulator of eCB-dependent plasticity. Interestingly, this applied equally to distinct forms of corticostriatal plasticity induced by either rate-based (LFS) or spike-timing-based (STDP) paradigm. We unveiled a novel mechanism by which BDNF shapes corticostriatal plasticity map. The corticostriatal axis is subjected to various forms of synaptic plasticity (Di Filippo et al., 2009; Lovinger, 2010). Long-term corticostriatal plasticity provides a fundamental mechanism by which the basal ganglia encode action selection, goal directed behavior and habit formation (Yin et al., 2009; Koralek et al., 2012; Shan et al., 2014; Rothwell et al., 2015; Hawes et al., 2015; Xiong et al., 2015; Ma et al., 2018; Perrin and Venance, 2018). Importantly, the retrogradely acting eCBs have emerged as major players in learning given their powerful influence in regulating corticostriatal plasticity (Lovinger, 2010; Mathur and Lovinger, 2012; Castillo et al., 2012; Araque et al., 2017). Synaptic plasticity is tightly controlled by various neuromodulators, also called the third factor (Foncelle et al., 2018), among which, BDNF appears to be indispensable for striatal functions (Baydyuk et al., 2011; Besusso et al., 2013; Unterwald et al., 2013). Indeed, in addition to its prominent role in inducing neuronal proliferation and differentiation, migration and survival, BDNF is a key regulator of synaptic transmission and plasticity in the adult brain (Carvalho et al., 2008; Waterhouse and Xu, 2009; Edelmann et al., 2014; Park et al., 2014). Intriguingly, compared to other neuronal networks where BDNF can act on both pre-and post-synaptic sites (Mohajerani et al., 2007; Sivakumaran et al, 2009; Inagaki et al., 2008; Edelmann et al., 2014), at the corticostriatal synapse BDNF is mainly released anterogradely and acts on postsynaptic TrkB (Baquet et al., 2004; Jia et al., 2010; Park et al., 2014). Such peculiarity is also supported by the low contents of BDNF mRNA and the high levels of TrkB in MSNs (Altar et al., 1997; Conner et al., 1997; Freeman et al., 2003; Fumagalli et al., 2007; Besusso et al., 2013). Thus, eCBs and BDNF follow opposite ways of action in striatum, i.e. retrograde and anterograde, respectively. Nevertheless, these two systems establish cross-talk in various central structures (Khaspekov et al., 2004; Huang et al., 2008; Maison et al., 2009; Lemtiri-Chlieh and Levine 2010; Roloff et al., 2010; Luongo et al., 2014; Zhao et al. 2015; Zhong et al., 2015; Bennett et al., 2017; Yeh et al., 2017; Maglio et al., 2018). This is exemplified by the IPSC depression in layer 2/3 pyramidal neurons induced by BDNF-generated CB_1_R activation (Lemtiri-Chlieh and Levine, 2010; Zhao and Levine, 2014). Such effect involves an increase of PLC-dependent eCB release from pyramidal neurons, which ultimately leads to a decrease of GABA release from presynaptic inhibitory terminals (Lemtiri-Chlieh and Levine, 2010; Zhao and Levine, 2014) and results in inhibitory LTD (Zhao et al., 2015). Despite the characterization of multiple cross-talk between BDNF/TrkB and eCB systems (Bennett et al., 2017), the role of BDNF in eCB-mediated long-term plasticity has not been determined yet. We thus questioned the impact of BDNF in the striatum where various forms of bidirectional eCB-mediated plasticity occur (Cui et al. 2016; Xu et al., 2018; for review see: Araque et al., 2017).

Information in the brain can be engrammed via two main activity patterns: the rate-and spike-time coding (deCharms and Zador, 2000), which can be mimicked experimentally by the use of two types of cell conditioning paradigms: rate-based and spike-timing-based protocols. Here, we show that BDNF regulates the expression of both rate- and spike-timing-based eCB-mediated long-term plasticity. Indeed, LFS-mediated LTD and bidirectional eCB-STDP are tightly dependent on the activation of TrkB. The common point of these LFS and STDP paradigms is that they were applied at 1 Hz presynaptic stimulation. This implies that BDNF might be efficiently released from cortical terminals at a low frequency, as recently observed in the hippocampus (Lu et al. 2014), and even for a low number of stimulations, as exemplified for the eCB-tLTP which requires only 10 pairings. Interestingly, a recent study has shown that eCBs set the inhibition strength onto pyramidal cells of the barrel cortex, and in turn to calcium spike facilitation and to a form of BDNF-dependent LTP (Maglio et al., 2018). This report, together with our own results, suggests that the involvement of BDNF in eCB-plasticity is a paramount mechanism at various central synapses.

Pyramidal cells from cortex layer 5 contact two MSN subpopulations belonging to the direct (striatonigral) or indirect (striato-pallido-subthalamo-nigral) striatal pathways (Gerfen and Surmeier, 2011; Calabresi et al., 2014). These two MSN subtypes express different dopaminergic receptors (D_1_R-like and D_2_R-like for the direct and indirect pathways, respectively). Since in the dorsal striatum, D_1_R- and D_2_R-expressing MSNs are roughly equally distributed (Gerfen and Surmeier, 2011; Gangarossa et al., 2013), we could expect that half of the MSNs would express heterogeneous eCB-plasticity and/or BDNF/TrkB regulation. We have previously shown that, depending on the activity patterns on either side of the synapse, eCB-tLTD and eCB-tLTP can be indifferently induced in both D_1_R- and D_2_R-MSNs (Paillé et al., 2013, Cui et al., 2015; Xu et al., 2018). In addition, we did not observe a dichotomy in plasticity occurrence as expected if D_1_R- or D_2_R-MSNs are differentially expressed eCB-plasticity. Importantly, cell-type deletion of TrkB in D_1_R-MSNs and D_2_R-MSNs has been associated to altered mRNA expression of GABA signaling (Koo et al., 2014) as well as to distinct phenotypes (Koo et a., 2014; Besusso et al., 2013). Although corticostriatal TrkB-eCB-plasticity occurs in most of the randomly recorded MSNs, these studies point to a specific role of TrkB in D_1_R-MSNs and D_2_R-MSNs. BDNF may contribute to corticostriatal events via its capacity to evoke presynaptic glutamate release (Zhang et al., 2013; Jovanovic et al., 2000; Yano et al., 2006; Park et al., 2014). However, in our experiments we did not observe significant modifications of EPSP amplitude following bath-application of DHF (TrkB agonist), or ANA12 and K252a (TrkB antagonists) (see Methods). This may be due to absence of GABA antagonists in our recordings to preserve the bidirectional and anti-Hebbian corticostriatal STDP (Paillé et al., 2013; Valtcheva et al., 2017); GABA antagonists are classically used to isolate glutamate and GABA events. It may then be possible that endogenous levels of striatal GABA may account for such apparent discrepancy.

BDNF is critical for neuronal maturation and striatal GABAergic neurons considerably mature between P_8_ and P_19_ (Santhakumar et al., 2010). STDP acquires bidirectional anti-Hebbian features in juvenile (P_20-25_), pre-adult (P_25-35_) and adult (P_60-90_) rat brains (Valtcheva et al., 2017) because of the tonic GABA component arising from P_16_ (Ade et al., 2008; Kirmse et al., 2008; Santhakumar et al., 2010). Since neurotrophins (NT3, BDNF and NGF) play a critical role in neuronal development and maturation, we used pre-adult (P_25-35_) brain slices in which BDNF mRNA is high and stably throughout adulthood (Maisonpierre et al., 1990; Timmusk et al, 1994). However, whether aging processes may impact BDNF-dependent corticostriatal eCB-plasticity remain to be established.

The effects of BDNF on synaptic plasticity are known to proceed via three major signaling pathways: PLCγ, MEK/ERK and PI3K/Akt (Patapoutian and Reichardt, 2001; Park and Poo, 2012; Edelmann et al, 2014). On the one hand, TrkB activation leads to the activation of PLCγ, which increases the levels of DAG and IP_3_. In turn this is expected to increase the production of 2-AG from DAG and to boost cytoplasmic calcium transients via IP_3_-dependent calcium release from internal stores. On the other hand, TrkB activates the MAPK/ERK and PI3K-Akt pathways, modulating plasticity through a range of potential mechanisms including AMPAR or NMDAR activity regulation. The PLCγ pathway, and the associated DAG and calcium boost upon TrkB activation, appears to be the best candidate for directly promoting eCB synthesis and release. It is well established that eCBs mobilization is promoted through transient increases in intracellular Ca^2+^ and activation of Gq-coupled receptors and subsequent PLC-dependent increase in DAG (Hashimotodani et al., 2007; Piomelli et al., 2007; Kano et al., 2009; Castillo, et al., 2012). BDNF is known to trigger local and fast transient increases in intracellular Ca^2+^ concentrations (Lang et al., 2007), an effect that has been attributed to slow mobilization of internal Ca^2+^ stores (Reichardt 2006; Amaral and Pozzo-Miller 2007) as well as to fast opening of voltage-gated Ca^2+^ channels (Kovalchuk et al., 2002). BDNF evokes transient calcium events in spines and dendrites of hippocampal granule cells and LTP is observed when these events are paired with weak synaptic stimulation (Kovalchuk et al., 2002). Using two-photon imaging combined with patch-clamp recordings, we show that TrkB activation modifies the kinetics of Ca^2+^ transients triggered by bAPs only, pre-post or post-pre STDP pairings. Indeed, TrkB activation promotes longer Ca^2+^ transients (this effect was prevented by TrkB antagonist), yielding a larger quantity of intracellular Ca^2+^ and potentially a larger eCB synthesis and release. Interestingly, we observed these modifications of Ca^2+^ transients in spines but not in dendritic shafts. Because of electrical compartmentalization, dendritic spines and shafts experience different voltage variations for incoming EPSPs, whereas it may not be the case for bAPs (Yuste, 2013). Synaptic activation triggers Ca^2+^-evoked events in spines, which propagate only partially in dendrites. Thus, for paired stimulation, we observed larger Ca^2+^-evoked events and more prominent effect of amplification by TrkB activation in spines compared to shafts.

To determine which pathway was required for TrkB-induced eCB-STDP (PLCγ, MAPK/ERK and/or PI3K-Akt pathways), we assembled a mathematical model describing the temporal dynamics of the biochemical reactions implicated in corticostriatal synaptic plasticity. The model accounted for the PLCγ pathway, disregarding the MAPK and PI3K-Akt pathways. Supporting this option, we showed that when MAPK and PI3K-Akt pathways were blocked, eCB-STDP could still be observed (Fig. 6), demonstrating that MAPK and PI3K-Akt pathways are not required for eCB-mediated plasticity at corticostriatal synapses. Beyond the very good match with the experimental data, the model predicted that TrkB agonists would facilitate the expression of eCB-tLTP and to a lesser extent of eCB-tLTD. Experimental validation of this prediction is a strong indication that TrkB modulation of eCB-tLTP proceeds via PLCγ, not MAPK nor PI3K. Namely, it was possible to enlarge the domains of expression of eCB-tLTP and eCB-tLTD by acting directly on TrkB activation as predicted by the mathematical model. Indeed, for pairings that did not induce eCB-tLTP and eCB-tLTD in control conditions, i.e. 25 post-pre and 25 pre-post pairings, respectively, (Cui et al., 2015; Cui et al., 2016; Xu et al., 2018), the activation of TrkB allows a considerable remodeling of the domain of expression of eCB-plasticity: when TrkB was activated by DHF during the 25 post-pre pairings, an eCB-dependent tLTP could be induced. Note however that, when DHF was applied all along the recording (i.e. the 25 post-pre pairings plus the following hour of recording) no plasticity was observed. These results may indicate that a brief and transient activation of TrkB is required for tLTP, whereas prolonged activation of TrkB would lead to tLTD. This is in line with the mechanisms that promote eCB-tLTD and eCB-tLTP: moderate levels and prolonged releases of eCBs lead to tLTD, whereas brief releases of large eCB concentration yield tLTP (Cui et al., 2016; Cui et al., 2018; Xu et al., 2018). This indicates that the temporal activation of TrkB is essential not only for the expression of eCB-STDP but also for its expression domains, in a way similar to eCBs (Cui et al., 2016). It is not currently possible to specifically block PLCγ activity because of the absence of a selective inhibitor that would exclusively target the γ isoform leaving the others (mostly *β* and *δ*) unaffected. The use of unspecific PLC antagonists (such as the widely-used U-73122) would prevent the synthesis of eCB by inhibiting PLC*β*, a key actor in 2-AG production (Castillo et al., 2012; Hashimotodani et al., 2007). Therefore, it remains to directly target the PLCγ to fully demonstrate the link between TrkB activation and eCB synthesis and release. Nevertheless, we opted for the strategy of blocking the MAPK and PI3K pathways and examine whether eCB-plasticity would be affected. We found that blocking specifically both the MAPK and PI3K pathways does not abrogate TrkB-mediated eCB-tLTD and eCB-tLTP. Experimental evidence and model results therefore converge to strongly suggests that the modulation of eCB-STDP by TrkB specifically acts via the PLCγ pathway, not the MAPK nor PI3K.

In neurodegenerative diseases, such as Huntington disease, (Strand et al. 2007; Zuccato and Cattaneo 2009; Park 2018; Yu et al. 2018) and neurodevelopmental disorders (Autry and Monteggia 2012; Baydyuk and Xu 2014; Deinhardt and Chao 2014), BDNF release is strongly diminished. In the light of our results, this reduction of BDNF account for the reduction or the loss of synaptic plasticity, as observed in Huntington disease rodent models (Kolodziejczyk et al. 2014; Plotkin et al. 2014; Sepers et al. 2018). On the opposite, BDNF levels are increased in maladaptive disorders such as ethanol or cocaine addiction (Jeanblanc et al. 2009; Im et al. 2010; Lu et al. 2010; Bahi and Dreyer 2013) or neurogenetic disorders, such as DYT1 dystonia (Maltese et al., 2018). In these cases, some aberrant plasticity has been observed (Deinhardt and Chao 2014; Zlebnik and Cheer 2016; Maltese et al., 2018) in line with our results showing that an over-activation of TrkB triggers a remodeling and an enlargement of the domains of expression of Hebbian plasticity. For instance, in DYT1 dystonia, increased BDNF levels trigger an abnormal LTP (together with structural dendritic modifications) during critical developmental window in premature MSNs (Maltese et al., 2018). In conclusion our study unveils a novel synaptic mechanism by which BDNF/TrkB governs striatal functions, thus potentially paving the way to a better understanding of a range of diseases in which alteration or disruption of BDNF/TrkB signaling has been recognized as causative feature for learning and memory deficits.

## ACKNOWLEDGMENTS

We thank the L.V. lab members for helpful suggestions and critical comments. This work was supported by grants from Fondation Patrick Brou de Laurière, Fondation pour la recherche sur le Cerveau (FRC) and Rotary Clubs de France (Espoir en Tête), INRIA, INSERM, Collège de France and CNRS. G.G is supported by Fondation Patrick Brou de Laurière.

## Conflict of interest

the authors have no competing financial interest to declare.

