## Supplementary material for "BDNF controls bidirectional endocannabinoid-plasticity at corticostriatal synapses"

### CAPTIONS TO SUPPLEMENTARY FIGURES

**Supplementary Figure 1: Representative STDP experiments (related the main Figures 2 and 3).**

**(A)** Example of tLTD induced by 100 pre-post pairings ($\Delta t_{STDP}$=-13±0.1ms) (baseline: 159±5pA, decreased by 56.7%, to 69±3pA, one hour after pairings.). Bottom, time course of Ri (baseline: 104.3±0.38 MΩ and 50-60 min after pairings: 111±0.4MΩ; change of 5.9%). (**B**) Example of the lack of plasticity after 100 pre-post pairings ($\Delta t_{STDP}$=8±0.1ms) with K252a (200nM) (the mean baseline EPSC amplitude was 119.2±2.62 pA before pairings and a lack of plasticity was observed one hour after pairings: 117.6±2.99 pA). Bottom, time course of Ri (baseline: 213±1MΩ and 50-60 min after pairings: 209±2MΩ; change of 1.8%). (**C**) Example of the lack of plasticity after 100 pre-post pairings ($\Delta t_{STDP}$=13±0.1ms) in presence of ANA12 (10µM) (the mean baseline EPSC amplitude was 187.2±3.73 pA before pairings and was not significantly altered one hour after pairings, 184.4±3.32 pA). Bottom, time course of Ri (baseline: 107±1MΩ and 50-60 min after pairings: 109±0.5MΩ; change of 2%).

(**D**) Example of tLTP induced by 10 post-pre pairings ($\Delta t_{STDP}$=-11±0.1ms) (the mean baseline EPSC amplitude was 230±6pA before pairings and was increased by ~67% to 383±5pA one hour after pairings). Bottom, time course of Ri (baseline: 114±1MΩ and 50-60 min after pairings: 116±1MΩ; change of 1%). (**E**) Example of the lack of plasticity after 10 post-pre pairings ($\Delta t_{STDP}$=-12±0.2ms) with K252a (200nM) (the mean baseline EPSC amplitude was 147±5pA before pairings and a tLTD induction was observed one hour after 10 post-pre pairings: 106±3pA). Bottom, time course of Ri (baseline: 64±1 MΩ and 50-60 min after pairings: 59±1MΩ; change of 8%). **(F)** Example of the lack of plasticity after 10 post-pre pairings ($\Delta t_{STDP}$=-13±0.3ms) in presence of ANA12 (10µM) (the mean baseline EPSC amplitude was 198±4pA before pairings and was not significantly altered one hour after pairings, 186±4pA). Bottom, time course of Ri (baseline: 74±0.2MΩ and 50-60 min after pairings: 68±0.2MΩ; change of 8%).

Insets correspond to the average EPSC amplitude during baseline (1, black trace) and the last 10 min of recording after STDP pairings (2, grey trace). Statistics (student t-test, first vs last 10 min of recording): ** $p<0.01$; *** $p<0.001$; ns, not significant.

**Supplementary Figure 2: Calcium transients in dendritic spines remain unchanged upon ANA12 and DMSO applications (related the main Figure 4)**.

(**A**) ANA12 and ANA12 followed by DHF application did not change the time course of Ca2+ elevations triggered by somatic current injections eliciting two bAPs in dendritic spines and shafts. (**B**) Two consecutive applications of DMSO (DMSO1 and DMSO2, 0.04% final concentration), to mimic successive applications of DHF and DHF+ANA12, did not change the time course of Ca2+ elevations in dendritic spines and shafts, triggered by two bAPs. (**C**) ANA12 and ANA12 followed by DHF application did not change the time course of $Ca^{2+}$ elevations triggered by pre-post paired corticostriatal stimulations in dendritic spines and shafts. Error bars represent the SEM. ns, not significant.

**Supplementary Figure 3: Amplitudes of calcium evoked-events in dendritic spines and shafts.**

Amplitudes of $Ca^{2+}$ evoked-events (normalized to control) were not different upon DHF, ANA12 and DMSO bath-application, in dendritic spines and shafts, triggered by (**A**) two bAPs (post-stimulation only), or by (**B**) pre-post pairings, consisting in single evoked corticostriatal EPSP paired with two bAPs with $\Delta t$ ~+15-20 ms.

Error bars represent the SEM. ns: not significant.

**Supplementary Figure 4: Long-duration bath-applied DHF did not promote tLTP for 25 post-pre pairings.**

Averaged time-courses showing an overall absence of plasticity observed after 25 post-pre pairings with DHF bath-applied during the whole recording (2/5 cells showed tLTP). Error bars represent the SEM. Statistics (student t-test, first vs last 10 min of recording): ns, not significant.
