## Supplementary figures and images for "BDNF controls bidirectional endocannabinoid-plasticity at corticostriatal synapses"

### Supplementary Figure 1

#### Supplementary Figure 1

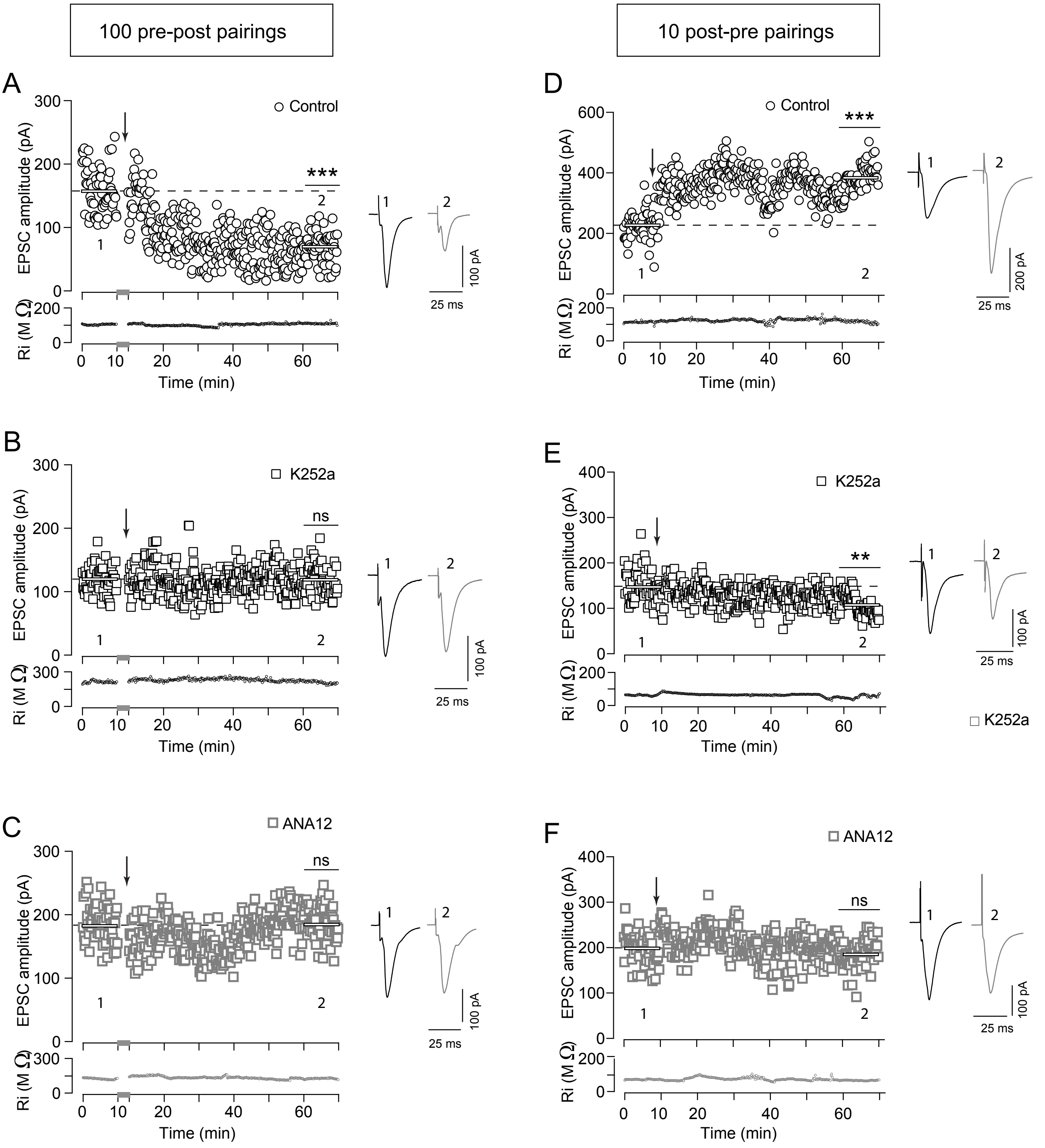

### Supplementary Figure 2

#### Supplementary Figure 2

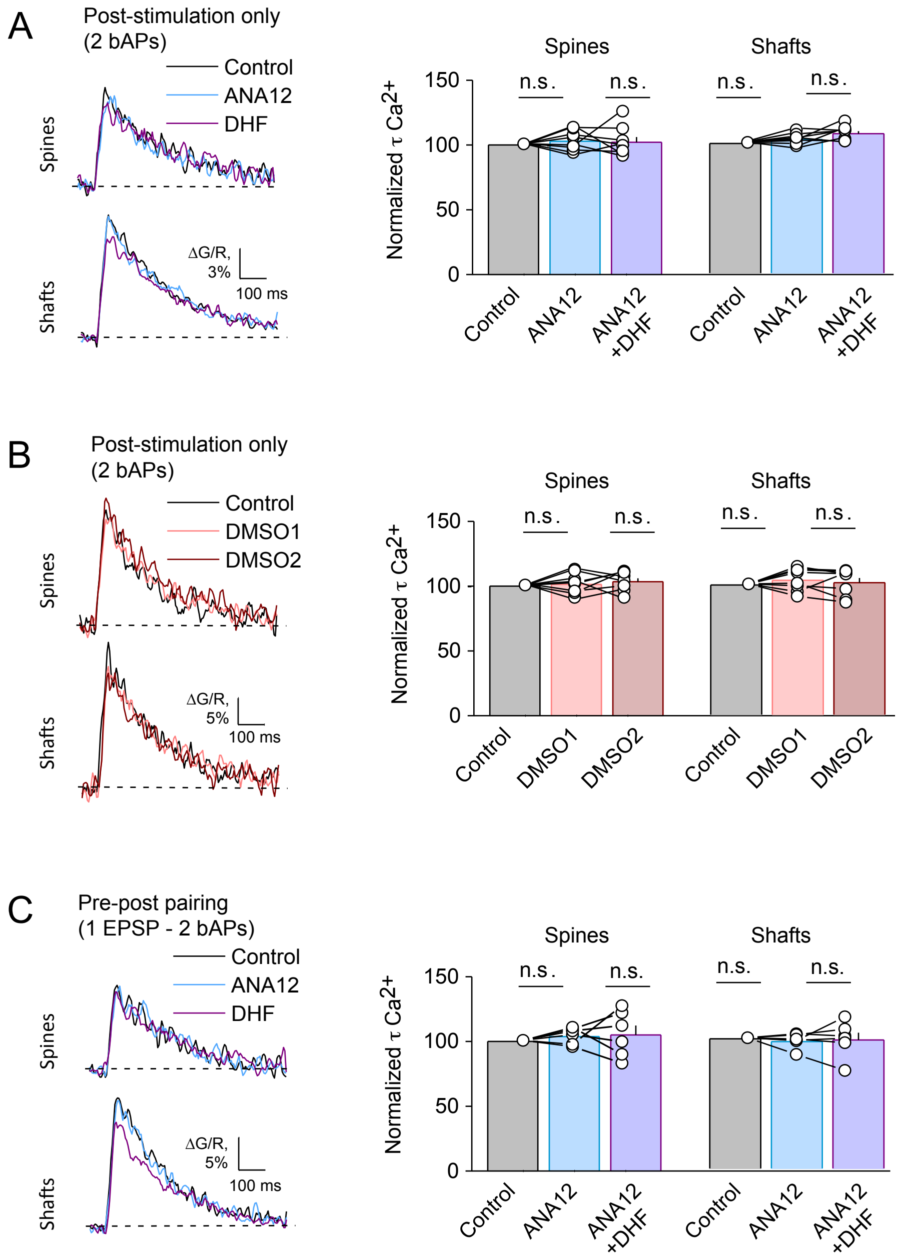

### Supplementary Figure 3

#### Supplementary Figure 3

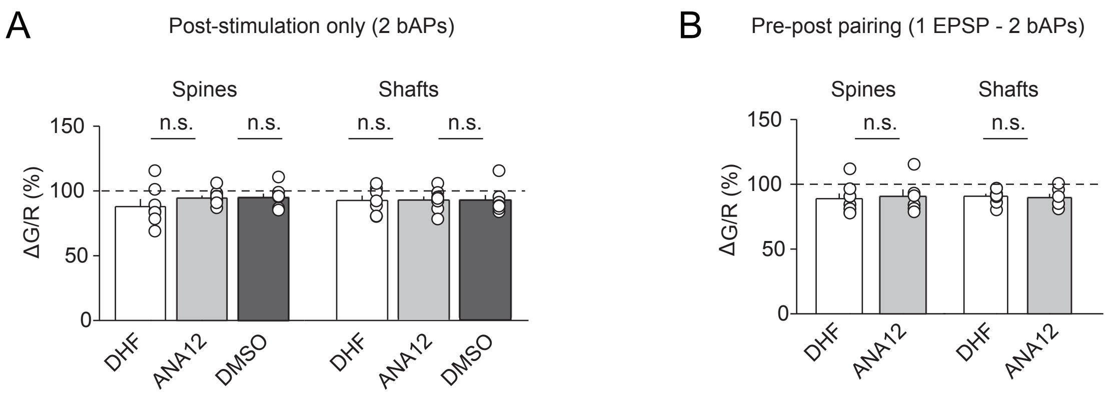

### Supplementary Figure 4

#### Supplementary Figure 4

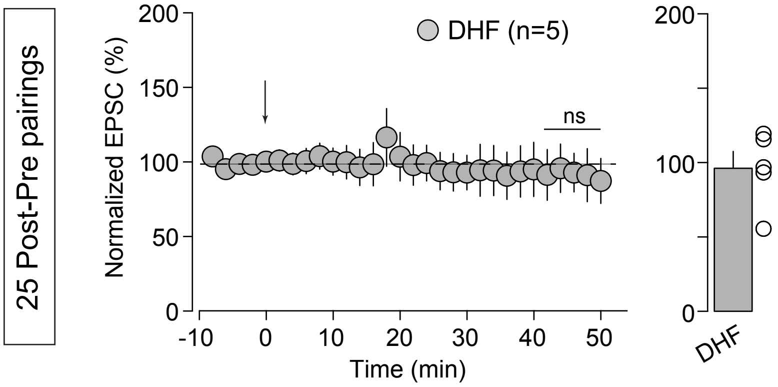
